# The human IG heavy chain constant gene locus is enriched for large structural variants and coding polymorphisms that vary among human populations

**DOI:** 10.1101/2025.02.12.634878

**Authors:** Uddalok Jana, Oscar L. Rodriguez, William Lees, Eric Engelbrecht, Zach Vanwinkle, Ayelet Peres, William S. Gibson, Kaitlyn Shields, Steven Schultze, Abdullah Dorgham, Matthew Emery, Gintaras Deikus, Robert Sebra, Evan E. Eichler, Gur Yaari, Melissa L. Smith, Corey T. Watson

**Affiliations:** Department of Biochemistry and Molecular Genetics, University of Louisville School of Medicine, Louisville, KY, USA; Department of Genetics and Genomic Sciences, Icahn School of Medicine at Mount Sinai, New York, NY, USA; Department of Microbiology, Icahn School of Medicine at Mount Sinai, New York, NY 10029, USA; Department of Genome Sciences, University of Washington School of Medicine, Seattle, WA, USA; Howard Hughes Medical Institute, University of Washington, Seattle, WA, USA; Bioengineering Program, Faculty of Engineering, Bar-Ilan University, Ramat Gan, 5290002, Israel; Vaccine Research Center, National Institute of Allergy and Infectious Disease, National Institute of Health, Bethesda, MD, USA

## Abstract

The human immunoglobulin heavy chain constant (IGHC) domain of antibodies (Ab) is responsible for effector functions critical to immunity. This domain is encoded by genes in the IGHC locus, where descriptions of genomic diversity remain incomplete. We utilized long-read sequencing to build an IGHC haplotype/variant catalog from 105 individuals of diverse ancestry. We discovered uncharacterized single nucleotide variants (SNV) and large structural variants (SVs, n=7), representing new genes and alleles enriched for non-synonymous substitutions, highlighting potential functional effects. Of the 221 identified IGHC alleles, 192 were novel. SNV, SV, and gene allele/genotype frequencies revealed population differentiation, including (i) hundreds of SNVs in African and East Asian populations exceeding a fixation index (F_ST_) of 0.3, and (ii) an IGHG4 haplotype carrying coding variants uniquely enriched in Asian populations. Our results illuminate missing signatures of IGHC diversity and establish a new foundation for investigating IGHC germline variation in Ab function and disease.

## Introduction

Antibodies (Abs) are critical components of the immune system^1^, playing pivotal roles in many disease states and clinical outcomes^2^. These proteins are produced by B lymphocytes and are either expressed at the cell surface as B cell receptors (BCRs) or are secreted as Abs. Abs and BCRs are composed of two pairs of identical ‘heavy’ and ‘light’ (kappa or lambda) chains, encoded by immunoglobulin (IG) genes located in three primary regions of the human genome: the IG heavy chain locus (IGH; chr14q32.33), and the light chain lambda (IGL; chr22q11.2) and kappa (IGK; chr2p11.2) loci. The expression of a given Ab depends on the somatic rearrangement of variable (V), diversity (D; heavy chain only), and joining (J) gene segments at IGH and one of the two light chain loci. The recombined V(D)J DNA template is then transcribed and spliced to a specific constant gene to comprise a full transcript. Genes within the IGH constant (IGHC) gene region encode and determine Ab isotypes (IgM, IgG, IgA, IgE, and IgD), which have multi-domain protein structures that contain binding sites for complement and Fc receptors^34^, and are responsible for modulating downstream effector functions, determining Ab transport, and Ab half-life^5^.

The IGHC genes reside within a ∼350 Kbp region adjacent to the IGHJ gene cluster^6^. Within the T2TCHM13v2.0 genome reference, for example, ten functional IGHC genes are present (IGHM, IGHD, IGHG3, IGHG1, IGHA1, IGHG2, two IGHG4 genes, IGHE, and IGHA2; listed in 5’-3’ order in Figure 1) each encoding a specific heavy chain isotype or subisotype. All IGHG genes share a common genomic structure consisting of CH1, CH2, and CH3 domain exons and a hinge region between the CH1 and CH2 domain^7^. The hinge exons of IGHG3 can vary in copy number, ranging from 2-4^8^. IGHD genes, not to be confused with D (“diversity”) gene segments that contribute to the antigen binding domain, have a structure similar to IGHG3, including two hinge region exons^9^. In contrast, IGHA, IGHE, and IGHM genes lack hinge region exons, and both IGHE and IGHM harbor an added CH4 domain exon downstream of CH3. The IGHC locus has been shaped by segmental duplication and consists of multiple homologous sequence blocks that harbor functional and pseudogenized paralogs of IGHG, IGHA, and IGHE genes^10,11^. The region is also diverse at the population-level, both with respect to structural variation that can result in changes in IGHC gene copy number^6,12–15^, as well as SNVs within IGHC coding sequences that result in extensive allelic variation^13,16,17^. Specifically, duplications and deletions involving IGHA1, IGHA2, IGHE, IGHG2, and IGHG4 have all been reported^6^, but the majority of these are yet to be characterized with nucleotide-level resolution. With respect to allelic variation, there are currently 100 alleles cataloged in the IMmunoGeneTics Information System database (date: November 24, 2022; IMGT; imgt.org;)^18^. However, recent reports indicate that there are likely to be many more alleles. A recent study of IGHG genes in a Brazilian cohort of 357 individuals from multiple population groups identified 28 novel alleles^16^; here, we use the term novel to refer to genes and alleles not curated in the International IMmunoGeneTics Information System database^19^. This was consistent with our recent study, in which we identified a total of 24 IGHG and 4 IGHM novel alleles in only 10 healthy individuals^13^. Furthermore, significant differences in IGHC allele frequency variation have been demonstrated between human populations^17,20^.

**Figure 1.**
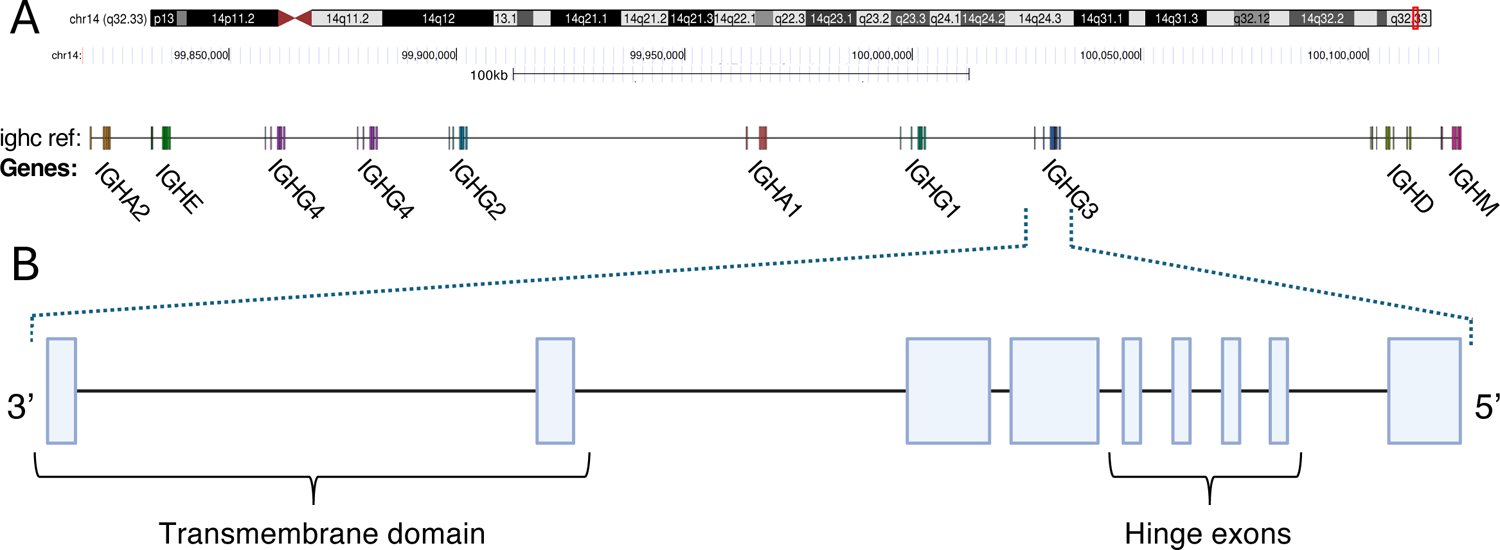
Schematic map of the immunoglobulin heavy chain constant (IGHC) region. **(A)** Genomic organization of the IGHC locus in T2T-CHM13v2.0 reference (chr14:99730306-100132055). **(B)** Expanded illustration depicts the exon-intron gene structure of IGHG3, representing the common structure of IGHC genes: three constant heavy (CH) domain exons CH1–CH2–CH3 (an additional CH4 domain is present in IGHM and IGHE), and hinge exons located between CH1 and CH2. Hinge exons vary by isotypes; 1-4 hinge exons occur in IGHG, IGHD but are absent in IGHA, IGHE, and IGHM. Depiction of 4 hinges as observed in IGHG3.

Functional consequences of IGHC polymorphism are known, but our understanding remains limited^21^. These functional consequences can include impacts on inter-individual variation in isotype distributions within the expressed Ab repertoire and serum Ab levels of various isotypes^14,22–25^. Additionally, coding allelic variation can alter protein function, such as induced effects on neonatal Fc receptor (FcRn) affinity^5^, and influence on Ab dependent cellular cytotoxicity (ADCC)^26^. These functional differences are expected to link IGHC polymorphism to complex disease phenotypes. Indeed, IGHG variants have been associated with disease outcomes in multiple contexts, including but not limited to multiple sclerosis (MS)^27^, human cytomegalovirus (HCMV) infection^28^, Myasthenia Gravis^29^, Myelin oligodendrocyte glycoprotein (MOG) antibody-associated disease (MOGAD)^30^, and COVID severity^31^.

Despite previous studies highlighting allelic diversity and structural variation within IGHC, our knowledge of locus-wide haplotype diversity at the genome-level remains insufficient. To date, there are only two fully curated and complete locus representations of IGHC. These include the T2T-CHM13v2.0 and GRCh38 reference assemblies, both of which are derived from individuals of Western European ancestry^32,33^. Historically, technical barriers have impeded the use of standard next-generation short-read sequencing to resolve complex loci^34,35^, specifically the IG loci^36–38^. The emergence of advanced long-read sequencing techniques, such as single molecule real-time (SMRT) sequencing (Pacific Biosciences), with enhanced accuracies, has shown significant improvements in comprehensive and precise detection of SNVs, SVs, repeat expansions, and complex genomic regions^35,39,40^, thereby providing a deeper and more accurate understanding of genomic architecture and variation. These technologies have the potential to enable higher-resolution characterization of the IGHC locus, including segmental duplications and deletions. Considering the anticipated extent of genetic diversity across human populations, in the short term, targeted approaches will be necessary to allow for high-throughput IGHC sequencing at population-scale.

In this study, we employ targeted long-read SMRT sequencing (Pacific Biosciences) of genomic DNA and large-insert fosmid clones from donors of the 1000 Genomes Project (1KGP) to generate high-quality haplotype resolved assemblies from five superpopulations (AFR, AMR, EUR, EAS, SAS). We first leverage a combination of fosmid clones and targeted long-read assemblies from four individuals of diverse genetic ancestries to generate manually curated and benchmarked high-quality haplotype-specific assemblies spanning the IGHC region. To facilitate additional benchmarking of our targeted long-read sequencing approach, we construct haplotype-phased assemblies for four trio probands, allowing for careful accuracy assessment utilizing parental datasets. We use these curated assemblies as the foundation for developing and evaluating a high-throughput computational pipeline for IGHC long-read assembly and variant calling. With this approach, we expand our survey of IGHC haplotype diversity to include an additional 97 individuals, leading to the largest curated set to date (n=105 donors). From this cohort, we identify extensive diversity at high resolution, including descriptions of novel SVs, SNVs, and short insertion-deletions (indels), as well as previously uncharacterized IGHC genes and alleles. This advancement enhances our understanding of IGHC gene diversity at the genome and population levels, highlighting vast and underappreciated polymorphism in this region. Our analyses illuminate the need to expand genetic surveys in these critical loci if we are to fully elucidate the importance of germline variation to Ab effector functions in disease.

## Results

### Building an initial IGHC haplotype reference set from orthogonal long-read datasets and parent-offspring trios

Given the paucity of available highly vetted haplotype assemblies for the complex IGHC region, we initially focused our effort on constructing high-quality haplotype-resolved assemblies from a selected number (n=4) of individuals representing diverse genetic ancestry. To do this, we followed our previous approach^41^, using orthogonal long-read SMRT sequencing data from target-enrichment capture libraries, and overlapping fosmid clones isolated from the same donors (Figure S1). For each donor, an average of ∼32 (range, 20-46) clones were selected, sequenced (RSII). and individually assembled. Through manual curation, overlapping clones were joined to create haplotype-resolved contigs. In addition, we constructed long-read IG-capture libraries^38,41,42^, generating HiFi reads (Sequel IIe) with average lengths of 5.4 Kbp, which were used to fill gaps and error correct haplotype-specific fosmid-based assemblies (Figure S1). Together, these data allowed us to construct 8 fully resolved gapless haplotype assemblies from NA18507 (AFR), NA18555 (EAS), NA19129 (AFR), and NA12156 (EUR) spanning the entire IGHC locus.

To construct additional phased reference assemblies, we conducted targeted long-read sequencing in 4 mother-father-child trios from the 1KGP: HG00671, HG00672, and HG00673 (EAS); HG01926, HG01927, and HG01928 (AMR); HG01950, HG01951, and HG01952 (AMR); and HG02715, HG02716, and HG02717 (AFR).

We generated IGHC assemblies using IGenotyper and Hifiasm from HiFi reads for all 12 individuals. Proband contigs were manually inspected alongside the parental assemblies to select the haplotype-specific contigs based on parent-of-origin (Figure S2). Selected contigs were merged to create 8 gapless haplotype-resolved IGHC assemblies. For 3 of the 4 probands, parental assemblies spanned >98% of their respective proband haplotypes. However, the maternally inherited haplotype of HG02717 was only partially covered due to haplotype drop-out in the maternal assembly (HG02716), likely due to a class-switch recombination event (Figure S3, Table S1).

We validated bases at each position in the hybrid and trio-based proband assemblies by alignment of HiFi reads for each sample back to their personalized IGHC reference. On average, we observed a mean of 41.5X coverage for each capture library relative to the respective haplotype-resolved assemblies. Across samples, we found that a median of 97.5% of haploid assembly bases had at least 10X coverage (Table S2). For these positions, we calculated the number of mapped reads supporting each base in the contig assembly, relative to the total number of reads mapping to that position. Requiring that at least 80% of mapped reads matched the base found in the assembly, HiFi reads supported >=99.9% of assembly bases (those with >9X coverage) across the 8 diploid assemblies (Table S2).

### Curating genetic diversity from high-quality IGHC haplotypes from diverse genetic backgrounds

From these 16 highly vetted haplotypes (Figure 2A), we curated variant call sets (SVs and SNVs), and IGHC genes and alleles. We detected 3 SVs (ranging in size from 188 bp to 19.5 Kbp) across the IGHC locus (Figure 2A). While earlier studies had noted the occurrence of IGHC SVs^43–45^, these represented the first curated descriptions of SVs with nucleotide resolution. The SVs identified among these 8 donors included: (1) an insertion-deletion (19.5 Kbp) variant region spanning the IGHG4 gene region, which was associated with variable IGHG4 copy number; (2) an intergenic inversion (4.5 Kbp) between IGHG2 and IGHGA1; and (3) an insertion-deletion (indel, 188 bp) variant that resulted in variable copies (3-4) of the IGHG3 hinge exons (Figure 2A). Focusing first on SNVs in gene regions, we next curated IGHC alleles from the 16 haplotypes. Consistent with recent reports^13,16,17^, we uncovered an extensive number of IGHC alleles, including those that were either incomplete or missing from existing databases (i.e., IMGT). In total, across the 10 annotated IGHC genes, we identified 70 alleles, 77% of which were not fully curated in IMGT (Figure 2B); alleles occurring at either of the two IGHG4 genes were consolidated into one set. Remarkably, no two haplotypes in this set of donors shared the same composition of SV and IGHC alleles (Figure 2A). Outside SV regions, we identified 3462 total SNVs across the 8 diploid assemblies. The number of SNVs varied between samples and was composed of both private SNVs (unique to a given sample) and those also observed in at least one other sample (Figure 2C). In general, we observed a greater overlap of SNVs between samples of the same genetic ancestry (Figure 2D). The majority of SNVs resided within intergenic regions, followed by those in IGHC exons and introns (Figure 2C). Across the 8 individuals, 7.25% (n=251) and 78% (n=2671) of SNVs were not found in either dbSNP or the 1KGP call sets, respectively (Figure 2E, F), a likely reflection of the difficulty in using short reads to detect IGHC variants. This was further demonstrated when we directly compared our SNV callsets to those generated by the 1KGP Phase 3 sets at the individual level, which revealed detected SNVs that were unique to either our long-read assemblies (∼76.4%) or the 1KGP (1.3%) callsets, indicating the likely detection of both false-negative and -positive genotypes called by the 1KGP using short reads (Figure 2G). This is consistent with results we have reported previously for variant calling in the IG loci^38,41,42^.

**Figure 2.**
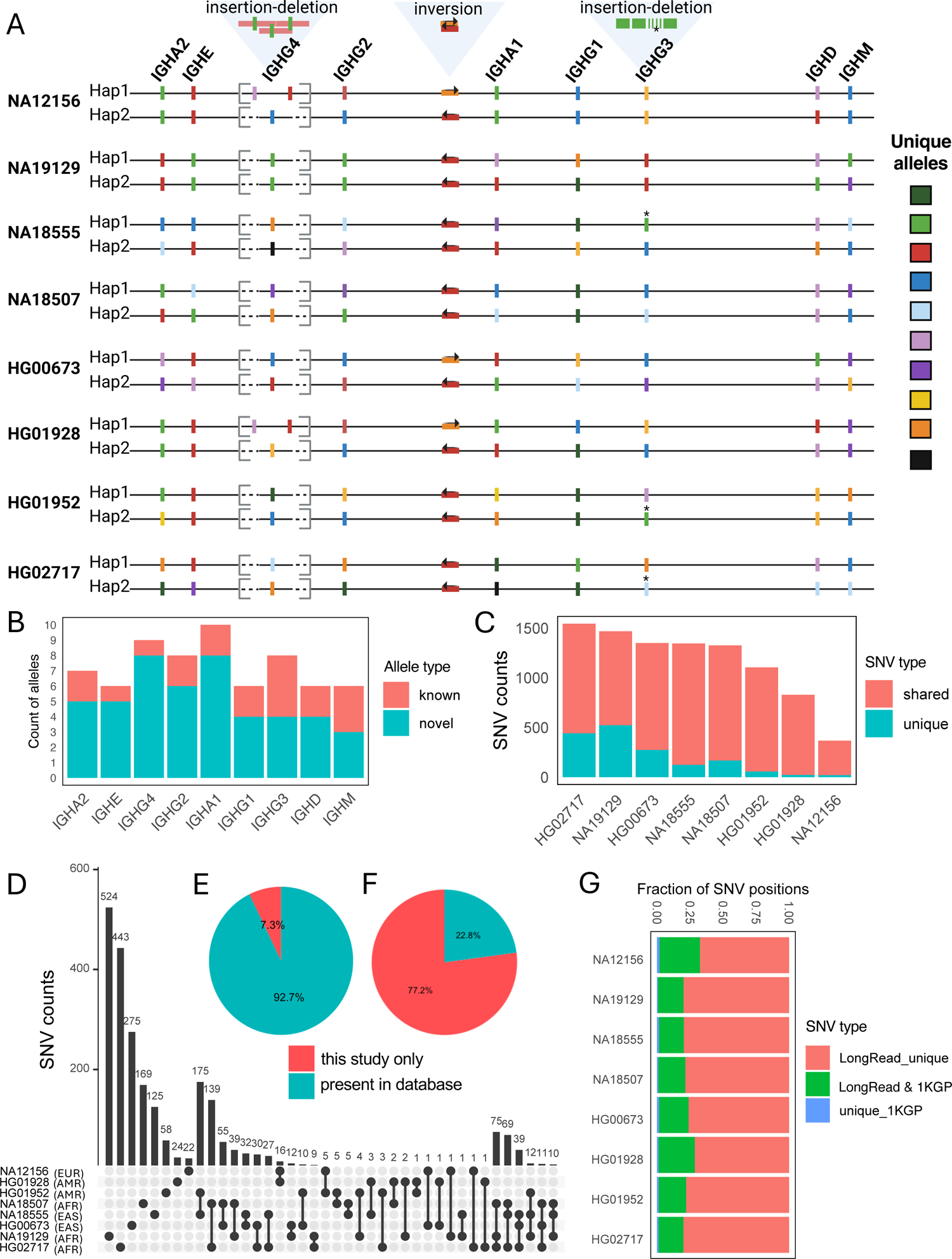
Resolving the IGHC locus in fosmid/trio-supported individuals from diverse ancestries. **(A)** Haplotype-resolved maps of the IGHC locus across individuals from diverse ancestries (n=8), indicating the locations of polymorphic genes (allelic variation denoted by color) and SVs. **(B)** Distribution of novel and known alleles (i.e., those present in IMGT) observed in the study cohort. **(C)** Counts of catalogued SNVs in each individual, partitioned by whether the SNVs are shared or private to each individual. **(D)** Upset plot showing the overlap of SNVs among individuals. **(E-F)** Comparison of observed SNVs to those curated in dbSNP_v155 (E) and the 1KGP-Phase-3 variant callset (F). **(G)** Comparison of individual SNVs from long-read sequencing (this study) and short-read data (1KGP-Phase-3).

### Assessing the performance of high-throughput IGHC targeted long-read sequencing in diploid samples

We next assessed targeted long-read sequencing data alone for generating accurate SNV/SV genotypes and gene/allele calls from haploid-resolved assemblies.

Prior to testing lab-generated data, we first evaluated the feasibility of our assembly approach using synthetic reads simulated from a subset of our high-quality IGHC diploid assemblies. We selected two resolved diploid benchmark assemblies along with the IGHC reference haplotype. For each genome, we simulated random reads of varying lengths (350 bp, 3 kb, 6 kb, and 10 kb), limiting coverage to 15X per haplotype. For the two diploid samples, reads from both haplotypes were merged into a single FASTA file and assembled using Hifiasm^46^. The resulting de novo assemblies were compared to the ground truth haplotypes by calculating average error rates per 10 kb window, demonstrating overall accuracy exceeding 99.9% for read lengths above 5 kb (Supplemental Table S3). This established clear expectations for assessing the performance of the experimental data generated from our benchmarking samples; specifically, read lengths generated by our protocol could be expected to yield accurate assemblies.

We next evaluated capture read coverage across all 8 diploid assemblies. For each haplotype, a mean of 95% of the bases were represented by >4X HiFi read coverage (Figure 3A). We benchmarked assemblies in these same donors using only targeted long-read data, processing the data through two pipelines, IGenotyper^38^ and Hifiasm^46^. IGenotyper adopts a reference-guided assembly strategy, using reads spanning phased blocks demarcated by haplotype-specific SNVs to generate haplotype-specific local assemblies from locally phased reads with Canu^47^. Hifiasm, in contrast, generates *de novo* diploid assemblies. On average, we found that IGenotyper yielded 27 IGHC assembly contigs per donor on average, with a mean length of 31 Kbp (Figure 3B). In contrast, Hifiasm generated 9 contigs per donor on average, with a mean contig length of 114.8 Kbp (Figure 3B). These results indicated that Hifiasm assemblies resulted in longer contigs across the locus overall. However, a modest number of phase-switch errors (in which a single contig has both paternally and maternally inherited bases) were observed in Hifiasm contigs in 4/8 individuals (Figure 3B). We also observed instances in which Hifiasm contigs included assembly errors caused by reads representing class switch recombination (CSR) events (Figure S4); however, this issue would not be expected to arise in non-LCL samples.

**Figure 3.**
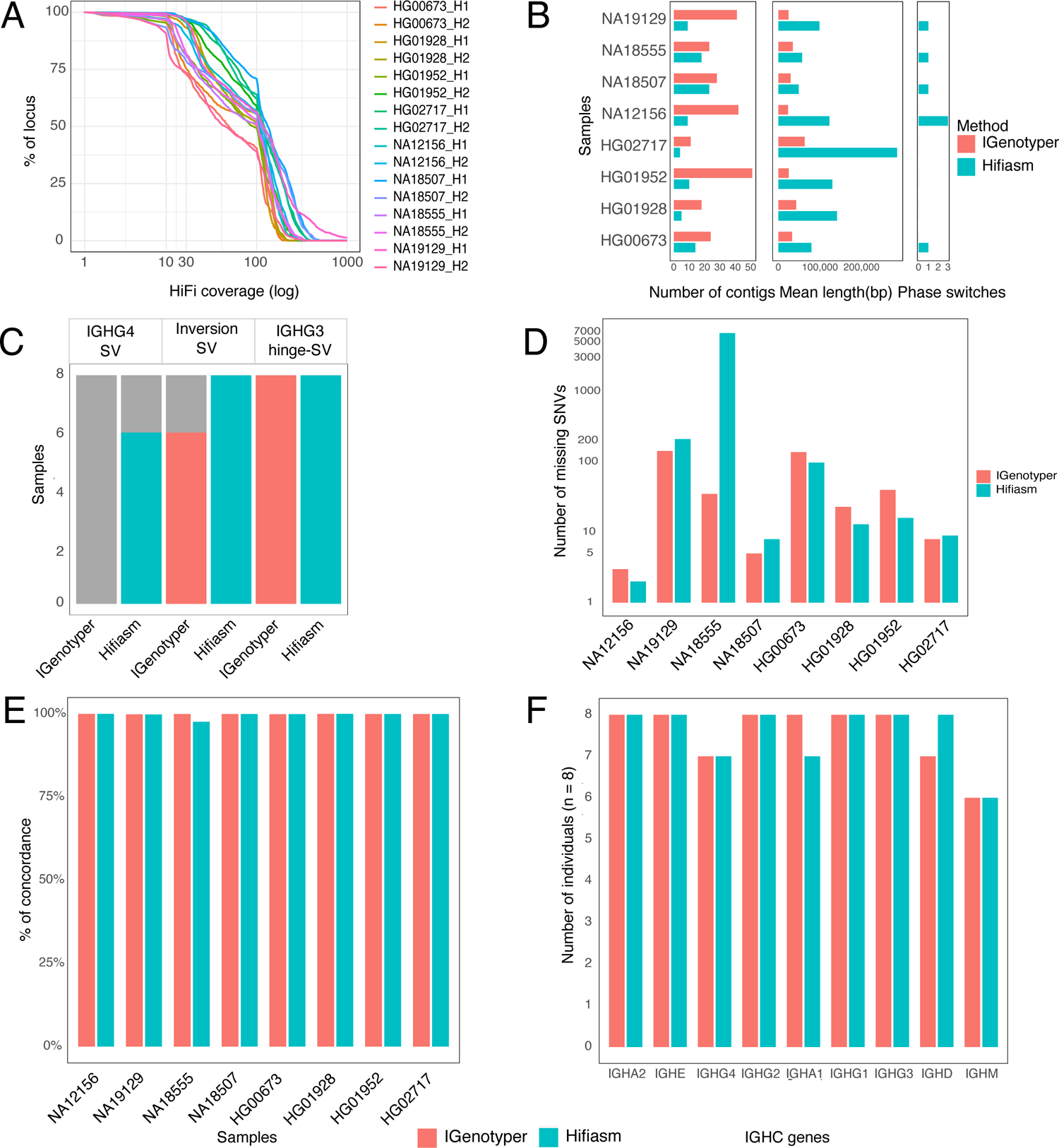
Benchmarking IGHC targeted long-read sequencing against diverse ancestral ground-truth assemblies. **(A)** Per-sample haplotype-specific HiFi read coverage: IGHC bases covered (Y-axis) vs read depth (X-axis). **(B)** Metrics (contig count, mean contig length, and number of phase-switch errors; see also Figure S4) for IGenotyper and Hifiasm pipeline generated assemblies. **(C)** Reproducibility of ground-truth common SV-annotations using automated IGenotyper and Hifiasm pipelines (required manual curation marked in grey). **(D)** Bar plots of variants in ground-truth assemblies missed by IGenotyper and Hifiasm assemblies. **(E)** Base concordance (%) between IGenotyper or Hifiasm assemblies relative to covered ground-truth bases. **(F)** Accuracy of alleles from IGenotyper and Hifiasm assemblies compared to the ground-truth allele set.

We next sought to assess the accuracy of SV and SNV calls generated by the IGenotyper and Hifiasm pipelines compared to ground truth hybrid/trio assembly-based call sets. We observed overall greater concordance between Hifiasm SV calls and ground truth sets, relative to those produced by IGenotyper (Figure 3C). For example, IGenotyper did not fully and accurately resolve IGHG4 SV genotypes in any sample. This was due to fragmented assemblies generated by IGenotyper over the IGHG4 region, which required manual joining for the curation of IGHG4 single- and 2-copy gene haplotypes in all samples. Likewise, while both Hifiasm and IGenotyper had perfect concordance with ground truth genotype calls for the IGHG3 hinge exon deletion, only Hifiasm accurately called the inversion SV genotype in all samples. Again, fragmented IGenotyper assemblies required manual curation to resolve inversion genotypes in a subset of samples.

To assess SNV call concordance, we considered genotypes at every base in each diploid assembly, whether per base genotypes were homozygous reference, heterozygous, or homozygous for the alternate non-reference allele. We assessed concordance between ground truth assemblies with IGenotyper and Hifiasm pipeline genotypes at each IGHC region base in the reference assembly. After excluding indels and SV-associated bases, the number of IGHC reference positions for which genotypes were called in the ground-truth datasets for each sample ranged from 288,503 to 288,985 base pairs. For each pipeline, genotypes were only called at a given position if spanned by a diploid assembly and a minimum of 4 HiFi reads. Positions not meeting both criteria were classified as “missing”. SNVs were called from the assemblies directly and mapped reads; in instances in which assembly-based SNV calls and read-based SNV calls were discordant, read-based genotypes were used. Genotype call sets from each pipeline were compared to those in the ground-truth assemblies. For IGenotyper call sets, the number of “missing” SNVs ranged from 3 to 143 (0.001% to 0.049%)(Figure 3D); for Hifiasm, the range was from 2 to 6761 (0.0006% to 2.34%)(Figure 3D). The sample with the high number of missing SNVs (n=6761) was caused by contig assembly dropout in this region, despite the occurrence of reads (Figure S5). Of the remaining positions, concordance ranged from 99.87% to 99.99% for IGenotyper, and 97.6% to 99.99% for Hifiasm (Figure 3E, Table S4).

Finally, we assessed IGHC allele calling accuracy. We developed a novel allele curation pipeline for extracting and annotating alleles for each IGHC gene present in either IGenotyper or Hifiasm assemblies, including an assessment of each annotation with respect to both assembly contig and HiFi read support (see STAR Methods). Using IGenotyper assemblies, the automated pipeline returned correct gene allele calls for all 8 donors over IGHA2, IGHE, IGHG2, IGHA1, IGHG1, and IGHG3 (Figure 3F). Accuracy was lower for IGHG4, IGHD, IGHM. For the IGHG4 miscalls, a split alignment over the IGHG4 SV region inhibited the automated extraction of genes from the haplotype-resolved contigs in a single individual; however, manual curation of these assemblies revealed the correct alleles. With respect to the IGHM and IGHD miscalls, we observed lower sequencing coverage over these genes in those donors, resulting in either the generation of partial contigs, or loss of contigs representing alternate haplotypes. However, in all cases, manual assembly-curation using the long-reads specific to each haplotype revealed that the correct alleles were present in all samples. Similarly, allele curation using Hifiasm assemblies achieved accurate assignments for all individuals across IGHA2, IGHE, IGHG2, IGHG1, IGHG3, and IGHD genes (Figure 3F), while correctly identifying alleles in 6 or 7 (depending on the gene) individuals for IGHG4, IGHA1, and IGHM. We suspect that the lower read coverage observed in these select samples reflected DNA loss associated with class-switch recombination represented in the LCLs.

### Assembling the IGHC locus using long reads in 97 individuals across 5 super-populations

To further characterize IGHC diversity we used IG-capture to conduct targeted long-read sequencing in 97 additional individuals from the 1KGP, increasing our total sample set to 105 individuals (AFR:n=16, AMR:n=28, EAS:n=22, EUR:n=19, SAS:n=20), encompassing 210 haplotypes. The mean read coverage for the IGHC reference locus across samples was 79.89X, ranging from 33X-250X. We processed all individuals through the IGenotyper and Hifiasm pipelines, to produce both reference-guided and *de novo* assemblies. The IGenotyper pipeline produced a mean of 27 contigs per individual with a mean length of 29.76 Kbp. Consistent with our benchmarking analysis (Figure 3), Hifiasm assemblies yielded an average of 12 contigs per sample, with an average length of 80 Kbp; in these samples, the IGHG4 segmental duplication/deletion region was fully resolved within a single contig for 208/210 haplotypes.

### Population-level genotyping allows for comprehensive characterization of additional structural variants, SNVs, and IGHC alleles

We first curated large SVs in our expanded cohort. Given the greater accuracy of Hifiasm assembly-based SV genotyping (Figure 3C), we used these assemblies for population-level assessment. As expected, we identified all three polymorphic SVs identified from the hybrid and trio-proband assemblies (Figure 2). In addition, we discovered the presence of 4 additional SVs, all of which were relatively rare in this cohort, occurring in <3 individuals. In two AFR individuals (HG03516, HG01891), we found evidence for a larger insertion variant, which was initially represented by the apparent presence of multiple distinct Hifiasm contigs overlapping the same region of the reference genome (Figure S6). However, manual curation of these contigs allowed for the reconstruction of a large insertion (∼120 Kbp), harboring an additional copy each of IGHE, IGHG4, IGHG2, and IGHA1 (Figure 4A). This SV was also recently discovered in an additional unrelated cohort using Oxford Nanopore Technologies (ONT)-based adaptive sampling sequencing^48^. Further, we identified a smaller insertion haplotype in one EUR individual (HG00096) that contained an additional copy of IGHG4, resulting in a tandem triplication of IGHG4 genes (Figure S7). In addition to these insertions, we also characterized multiple gene deletion events in EAS individuals. This included a deletion that resulted in the complete loss of both copies of the IGHG4 tandem duplication, which was observed in two EAS individuals (NA18543, NA19062); and a second larger deletion in a single EAS individual (NA18956) resulting in a putative fusion of the IGHA2-IGHA1 genes, and the loss of IGHE, IGHG4, and IGHG2 genes (Figure 4A, Figure S8).

**Figure 4.**
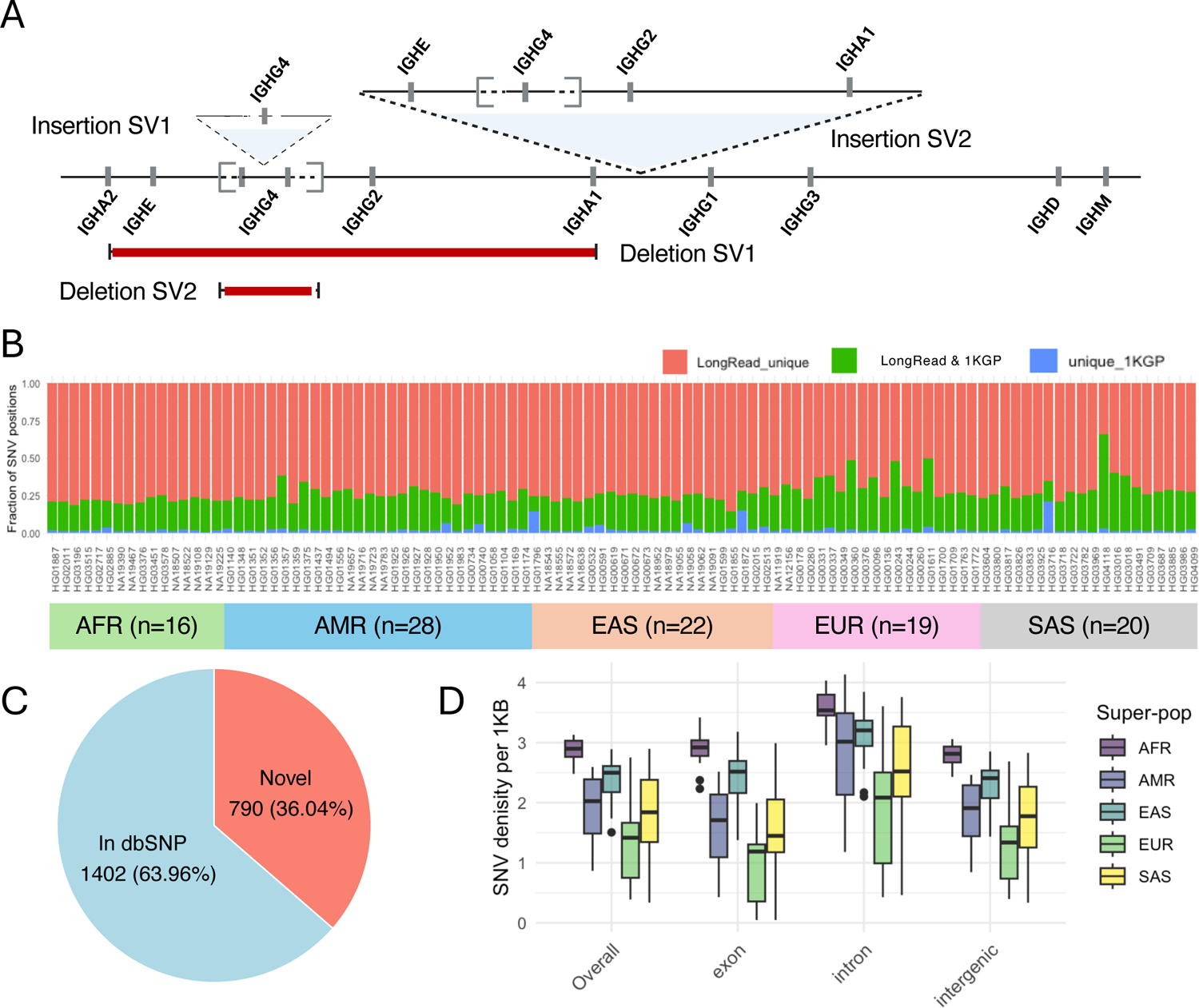
Characterization of SVs and SNVs from population-level dataset. **(A)** A map of additional characterized SVs in expanded population-level cohort: a tandem triplication of IGHG4 (Insertion-SV1, see also Figure S7); ∼120Kb insertion including paralogs of IGHE, IGHG4, IGHG2, IGHA1 (Insertion-SV2, see also Figure S6); large deletion spanning IGHE, IGHG4, and IGHG2 genes (Deletion-SV1, see also Figure S8), resulting putative fusion allele of IGHA1 and IGHA2 genes; deletion of IGHG4 genes (Deletion-SV2). **(B)** Per-sample SNVs identified from long-read datasets relative to 1KGP-Phase-3 SNV callsets. **(C)** Proportion of common SNVs (MAF >= 5%) in the expanded cohort present in dbSNP_v155 (see also Table S5, S6). **(D)** IGHC SNV density across samples in the expanded cohort based on genomic features and superpopulation.

After accounting for SVs, we next characterized SNVs across the IGHC locus. We identified a total of 5,889 variant sites; 92 of these sites were multi-allelic. Among these, 2,192 SNVs were observed at a minor allele frequency (MAF) ≥ 5%. Consistent with observations made in our benchmarking dataset, 790 (36%) of these common SNV sites were not represented in dbSNP (Figure 4C, Table S5). Additionally, sample-level comparison of SNV positions identified in our study and those derived from short-read data (1KGP) revealed that, on average, 73% of SNV positions were uniquely detected in our curated assemblies (Figure 4B, Table S6). This is consistent with the well-known limitations of short reads for accurately genotyping genomic regions enriched with duplicated sequences^49–51^. Together, these results highlight potential pitfalls of previous studies that leveraged array and short-read sequencing-based genotyping for disease association analysis^51^.

Alleles were annotated for all functional IGHC genes present in haplotypes from the 105 individuals. Because duplicate genes identified in the insertion haplotypes do not yet have official gene names, alleles were aggregated across all paralogs (i.e., IGHE, IGHG4, IGHG2, and IGHA1 duplicates) for simplicity. In total, we identified 221 alleles, only 29 of which (13%) were documented in the IMGT database (Figure 5A). Of the putative novel alleles, coding bases annotated from assemblies for 158 alleles were supported by at least 8 HiFi reads (see STAR Methods) in at least one individual (Figure S9); additionally, 91 of these alleles were also observed in >1 individual (Figure S9). Putative novel alleles were identified for all 9 IGHC genes, and outnumbered previously curated alleles for each gene. Notably, 71 alleles in the utilized IMGT reference set were not observed in our cohort, including alleles for IGHA2 (n=2), IGHE (n=3), IGHG4 (n=6), IGHG2 (n=15), IGHA1 (n=3), IGHG1 (n=12), IGHG3 (n=24), IGHD (n=3), and IGHM (n=3). The absence of these alleles in our data suggests they may represent rare variants or potential artifacts, akin to those noted for other IG loci^52^.

**Figure 5.**
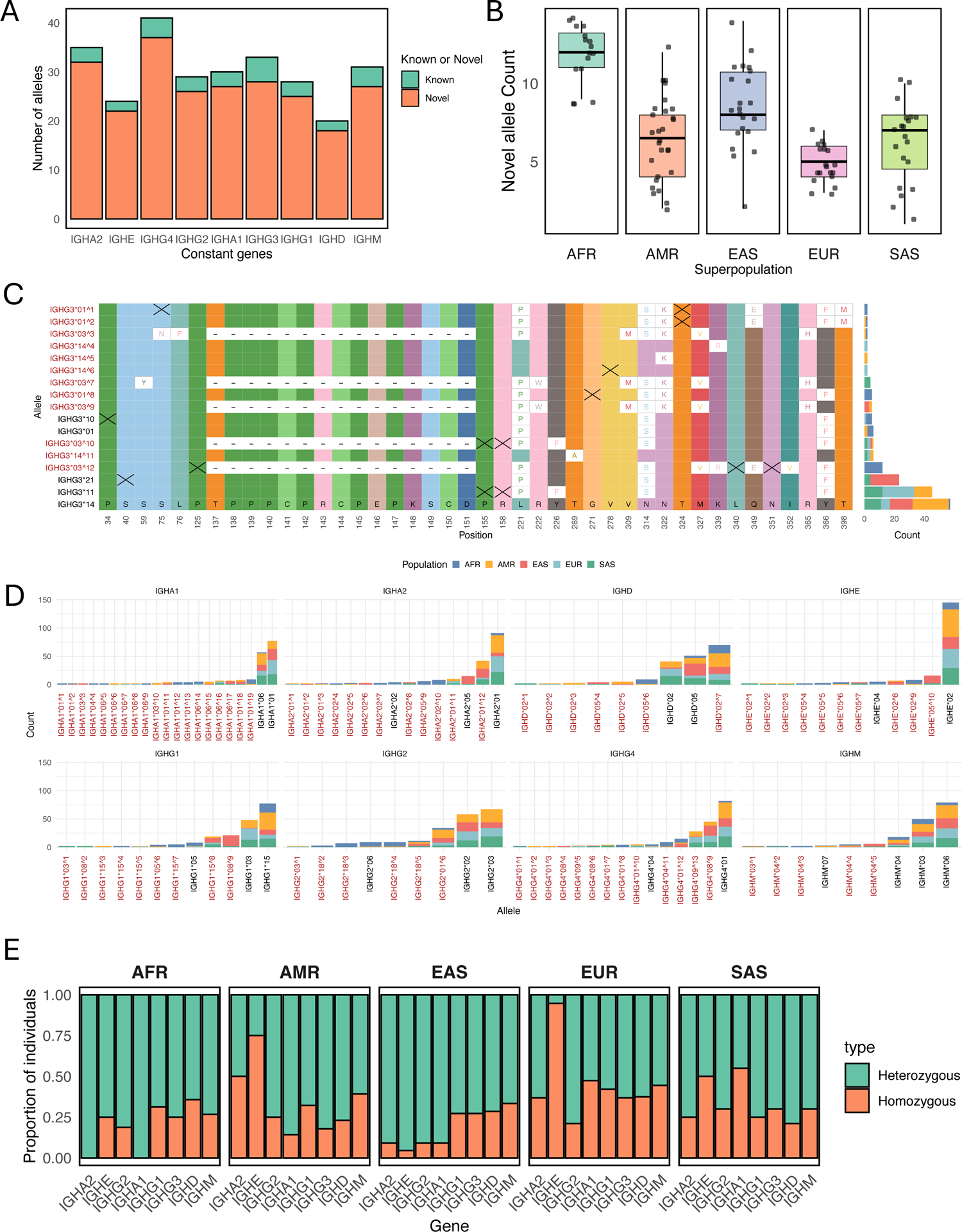
Characterization of IGHC allelic diversity from population-level dataset. **(A)** Counts of known and novel alleles for 9 IGHC genes; with close paralogs (e.g., IGHG4 duplicates) merged into a non-redundant set. **(B)** Boxplot showing novel allele distribution per individual across super populations (see also Figures S9, S10). **(C-D)** Multiple sequence alignment of IGHG3 alleles identified in our cohort (n=105); only variant positions are shown for alleles found in >1 individual, with “X” denoting synonymous variants. Stacked barplot shows allele counts by population (see also Figures S11, S12, S13). **(E)** Stacked bar plot showing gene wise heterozygosity/homozygosity, grouped by superpopulation.

All 105 individuals carried at least 1 novel allele, with individuals of African descent harboring ∼12 novel alleles on average (range: 8-14); this was in contrast to individuals of European descent, which were found to carry on average ∼5 novel alleles (range: 3-6)(Figure 5B; Figure S10). Well-supported novel alleles included both non-synonymous and synonymous nucleotide differences, as indicated in the example shown for IGHG3 alleles (Figure 5C; see also Figures S11, S12). The distributions of known and novel alleles varied widely among super populations, including a number of alleles observed in only a single population (Figure 5C and 5D). Not surprisingly, high heterozygosity rates for IGHC genes were observed across all populations (Figure 5E), with the majority of individuals carrying >1 allele per IGHC gene (Figure S10). A notable exception was the IGHE gene in both AMR and EUR individuals, which showed reduced heterozygosity relative to other populations and genes (Figure 5E).

The diversity discovered here raised questions concerning potential functional impacts. To assess this, we examined amino acid variation across all IGHC alleles in our cohort and compared these to known functional residues ^7,25,53^. None of the observed IGHA1 and IGHA2 alleles carried polymorphisms at known functional sites. In contrast, the IGHG genes showed substantial variability within previously reported functional residues (Table S7, S8). IGHG3 exhibited amino acid substitutions at 16 distinct residues, with at least one allele containing a variant at each curated functional site. These variants, as reported in previous studies^7,25,53^, have the potential to impact Fc-FcγR interactions, antibody-dependent cellular cytotoxicity (ADCC), and antibody half-life. IGHG1 and IGHG2 each harbored substitutions at four functional sites across observed alleles, while IGHG4 showed changes at two such residues. These positions have been associated with Fc-FcγR and FcRn interactions (Figure S13), suggested to impact downstream effector functions (Table S7, S8).

### SVs and IGHC allele frequencies vary between human populations

Mounting evidence indicates that signatures of haplotype diversity in the IG loci, including examples of SVs and both coding and non-coding SNVs, can differ by genetic ancestry ^16,17,20,54–56^. We leveraged the comprehensive IGHC SV, SNV, and allele genotype call sets generated here to identify differences between human populations represented in our cohort.

First, for all three common SVs, we observed significant variation between populations (Figure 6). For example, we observed that IGHG4 copy number was lower in EAS donors than all other populations (Figure 6A). Similarly, we observed that the inversion SV was invariant among donors of AFR populations, in contrast to all other populations in which both inversion alleles were found (Figure 6B). We also identified 14 haplotypes (∼7%) lacking a hinge exon in IGHG3 (Figure 6C), consistent with that previously reported^17^. These deletion haplotypes were observed in 12 individuals: two homozygotes (HG02011-AFR and HG03491-SAS), and 10 heterozygotes. Population-level analysis of the IGHG4 gene duplication suggested the potential of multiple recurrent deletions and duplications. Specifically, based on the mapping of single-copy IGHG4 spanning contigs to the reference assembly we identified several distinct putative breakpoints (Figure S14). This signature suggested the potential occurrence of more than one deletion and/or duplication event, as well as the potential for sequence divergence between SV haplotypes. Given this complexity, we concluded that the delineation of specific breakpoints relative to a single reference genome was fraught, and likely to lead to false inferences of breakpoints, SNV genotypes, and mis-assignment of single copy IGHG4 genes and alleles. Thus, to genotype this region, we employed a haplotype clustering approach, based on the pairwise distances of alignments of all IGHG4 spanning haplotypes (see STAR Methods). We identified 23 haplotype clusters, including 1 cluster representing 2-copy IGHG4 haplotypes (i.e., sharing the same structure as the reference assembly haplotype), 1 unique 3-copy haplotype, 2 unique haplotypes with a complete IGHG4 deletion (0-copies), and 19 single-copy IGHG4 haplotype-associated clusters. Cluster 1 (a single-copy cluster) was the most frequent across the cohort (n=74 haplotypes), followed by 2-copy IGHG4 haplotypes (n=30) and cluster 2 (n=22 haplotypes). Analysis of haplotype-associated alleles revealed that the most frequent clusters (1 and 2) alongside nine other clusters (5, 6, 9, 10, 11, 13, 15, 17, and 20) primarily harbored *01 and similar alleles (novel alleles with a closest match to *01), whereas clusters 7, 8, 12, 16, and 18 carried *04 and related alleles, and clusters 3, 4, 5, and 21 carried *08, *09 and related alleles (Figure 6F, Table S9).

**Figure 6.**
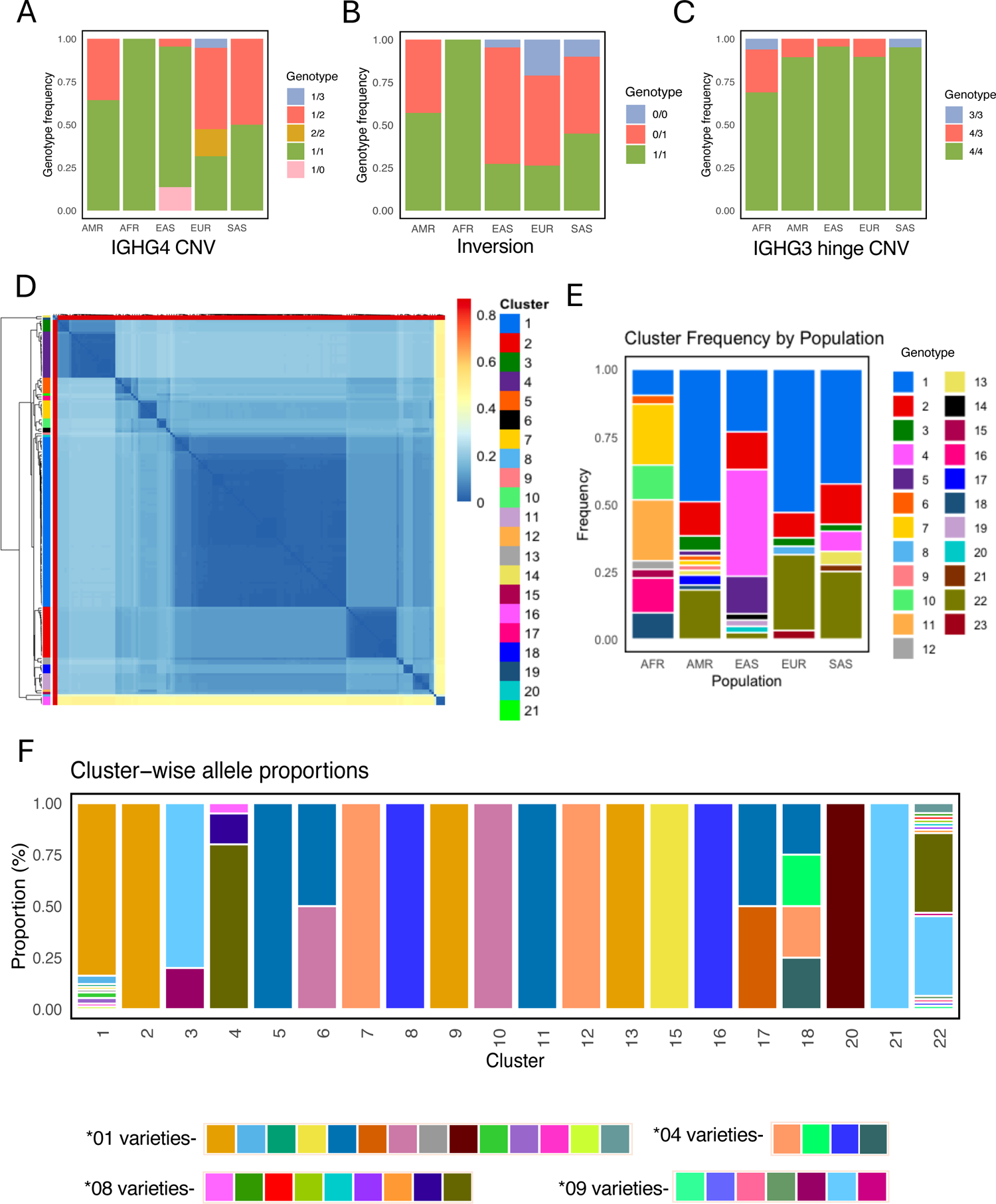
Comprehensive characterization of SVs across diverse genetic ancestries. **(A-C)** Stacked bar plot representing genotype frequencies by superpopulation for three common SVs. **(D)** Heatmap of pairwise nucleotide distances between IGHG4 region haplotypes; clustered hierarchically (dendrogram, left), with sequences in the heatmap are labeled by haplotype cluster, defined at 0.1 distance cutoff (see also Figure S14). **(E)** Stacked bar plot of IGHG4 haplotype cluster frequencies across super-populations **(F)** Stacked bar plot showing IGHG4 allele distribution across haplotype clusters.

Notably, when analyzed at the resolution of haplotype clusters, we found that the frequency of observed haplotypes varied considerably across populations (Figure 6E). Of single-copy haplotypes, clusters 7 and 11 were predominant among AFR individuals, whereas cluster 1 was the most common among EUR, AMR and SAS groups alongside the 2-copy IGHG4 haplotype (cluster 22). In addition, cluster 4 was observed in 17 out of 42 haplotypes in the EAS population. Interestingly, a greater number of clusters were observed in AFR and AMR individuals (9 and 11, respectively) whereas EUR had only 6 clusters. It is worth noting that validation of these observations will require larger cohorts. This will be particularly important to better understand population frequencies of the rarer SVs characterized here.

### Coding and non-coding SNVs highlight regions of IGHC exhibiting intra-population differences

A strength of our dataset is our accounting for the presence of SVs in our assemblies for improved SNV calling and genotyping. Thus, in addition to assessing population-level variation for SVs, we asked whether SNVs outside of SV regions also exhibited signatures of population divergence. We first assessed this by conducting principal component analysis (PCA) using common SNVs in non-SV regions; individuals with large, rare SVs (n=2, AFR; n=1, EAS) were excluded. Assessment of the first three principal components (PCs) indeed revealed separation by super-population (Figure 7A & B); in particular, EAS and AFR samples were the most differentiated along PC2 and PC1, respectively. We next tested SNV-level allele frequency differentiation using pairwise F_ST_ (Weir and Cockeram^57^), comparing each population group against all remaining groups collectively. Consistent with results from the PCA, drastically higher F_ST_ values were observed for AFR (F_ST_ as high as 0.95) and EAS (F_ST_ as high as 0.86). For three of the five population comparisons, we observed F_ST_ values that exceeded 0.3: 742 for AFR, 280 for EAS, and 74 for EUR. Interestingly, these high-F_ST_ (>0.3) sites were observed in IGHG1 and IGHG3 for all three of these populations (AFR, EAS and EUR). However, strong differentiation of AFR was also uniquely observed for sites in IGHM, IGHA1, and IGHG2 (Figure S15).

**Figure 7.**
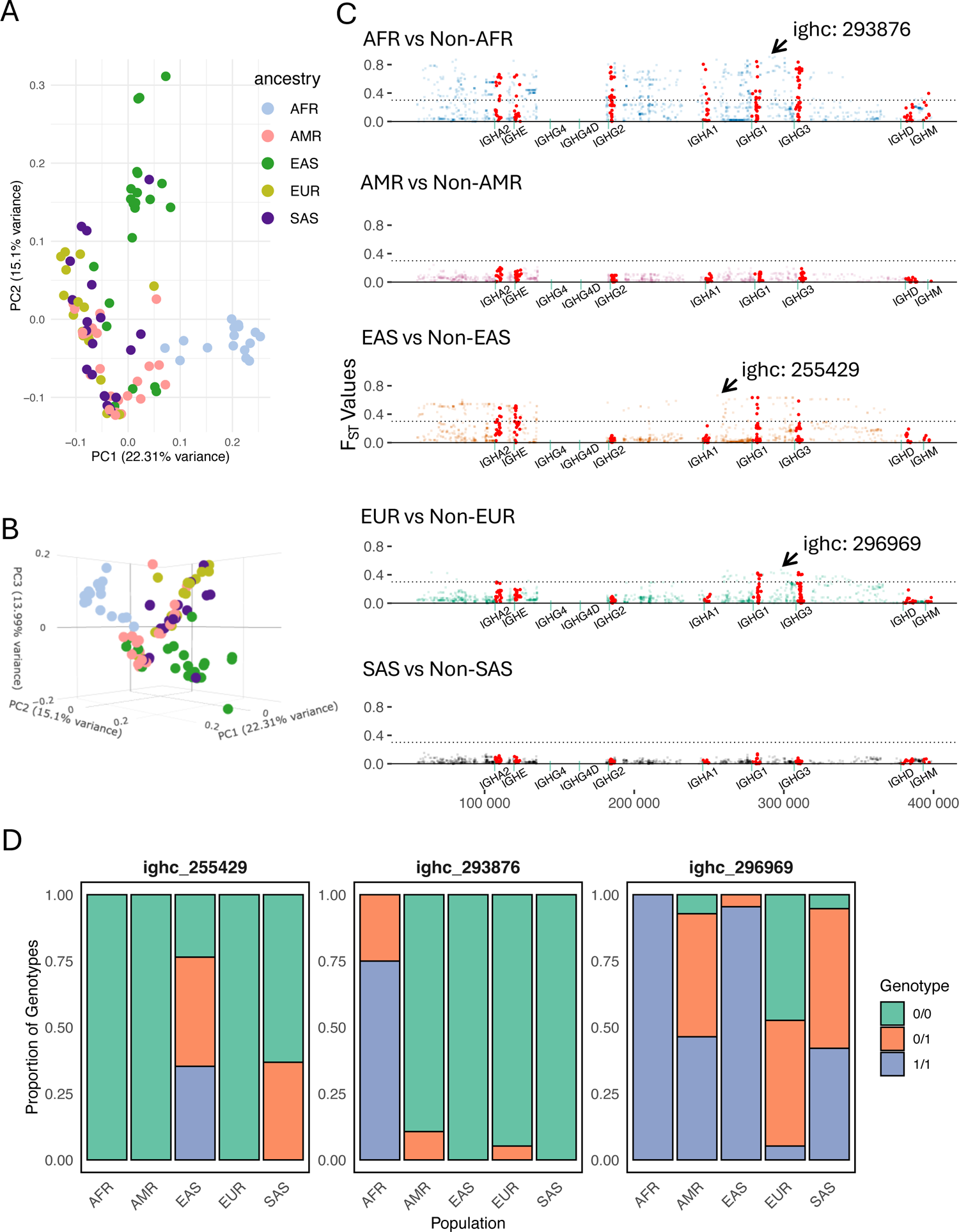
Coding and non-coding IGHC SNVs reveal inter-population genetic differences. **(A-B)** Principal component analysis of IGHC SNVs; points represent individuals colored by super-population. **(C)** Pairwise F_st_ values for IGHC SNVs comparing each population to all others (see also Figures S15, S17). **(D)** Genotype frequencies of three SNVs with highest F_st_ values across three populations.

## Discussion

IGHC genes encode critical domains of expressed Abs, facilitating downstream effector immune functions. Despite the biological importance of IGHC genes, their roles in health and disease, and relevance to therapeutic development, our understanding of IGHC genetic diversity has remained limited. Here, extending our previously published methods, we demonstrated the utility of a high-throughput molecular and informatics protocol that leverages highly accurate, long-read sequencing for reconstructing haplotype-resolved assemblies across the IGHC locus. Using an initial benchmarking set of vetted assemblies, we first showed that our pipeline provides accurate, locally phased assembly contigs representing all IGHC genes across the locus using a single molecular assay. Further, we demonstrated that our automated analysis workflow was able to generate reliable SNV and SV callsets and curate IGHC alleles (both novel and known). Second, by applying this approach at scale in the largest surveyed cohort to date, we characterized an extensive range of genomic variants, including undocumented SNVs and SVs, as well as novel IGHC genes and coding alleles, each with substantial inter-population variation. For the first time, these data provide a comprehensive snapshot of the tremendous IGHC diversity circulating in the human population.

The IGHC locus as represented in current linear reference genomes (i.e., GRCh38 and T2T) is composed of many large segmental duplications. It is now understood that these references are missing genes present across populations. This complex genomic architecture and “missing” data have thwarted previous efforts to accurately reconstruct IGHC haplotypes with nucleotide-level resolution using short-read sequencing. As a result, our knowledge of SVs in the IGHC region almost entirely draws from those inferred from RFLP assays, targeted PCR, and expressed Ab repertoire data ^6,14,15,44,58^. Thus, even though large polymorphic SVs in the locus have been indicated since the 1990s ^6,12,14,44,45^, the majority of these have remained undescribed and uncurated. Complete haplotype-resolved IGHC assemblies using a combination of Oxford Nanopore and PacBio HiFi data from multiple donors were recently curated, allowing for the first characterization of several large duplications, each harboring additional copies of IGHC genes ^48^. The largest of these included a duplication of ∼120 Kbp, spanning 4 functional genes. Critically, these duplicated genes, which represented both IgG and IgA subisotypes were shown to be expressed, highlighting deficits in our basic knowledge of IG gene regulation and fundamental Ab biology. Here, we drastically expanded on this survey of IGHC SVs, identifying 7 SVs in total across 210 haplotypes and five superpopulations. Importantly, our assembly resolution allowed us to delineate the presence of multiple overlapping SVs, indicating that the IGHC locus is a SV hotspot, and a site of recurrent events. With respect to the large multi-gene SV noted above, we characterized IGHC haplotypes from 2 individuals (HG01891, HG03516) carrying a previously described large segmental duplication and additionally resolved a multigene deletion haplotype in one EAS individual (NA18956) that resulted in the fusion of IGHA2 and IGHA1. In a second subregion of IGHC spanning IGHG4 and its close paralogs, we identified 23 distinct haplotype clusters, which were defined by variation in IGHG4 gene copy number (0-3 haploid copies), differences in nucleotide/SNV profiles, coding alleles, and variable putative SV breakpoints. Notably, IGHG4 was previously reported to represent a site of local population-specific adaptation and Neanderthal introgression^20^. In our cohort we observed that the frequency of IGHG4 haplotypes varied considerably between human populations, with the AFR population exhibiting the greatest diversity in terms of the number of unique clusters and their frequency distributions.

Our resolution of SVs also allowed us to improve the accuracy of SNV detection and genotyping. Specifically, haplotype-level characterization of SVs enhanced variant calling by reducing misclassification of paralogous sequence variants (PSVs) within closely related segmental duplications as SNVs. Demonstrating this, we compared the variant call set derived from our long-read sequencing data to that obtained by the 1KGP for these same samples. Interestingly, we found that a median of 74.3% of SNVs were only detected in our long-read-based variant call set. We also found that 36% of common (MAF ≥ 5%) SNVs in our dataset were not present in dbSNP. Notably, even at the individual level, we observed that on average, 73% of SNVs genotyped in our long-read data were not called by the 1KGP. These data indicate the additional work needed to better integrate missing variation in genetic association studies. The implications of these missing data for previously conducted GWAS is unknown but has likely negatively impacted the discovery of IGHC associations in disease. The same pitfalls that obfuscate SNV genotyping in complex loci can also impede accurate curation of genes and alleles. Our data clearly demonstrate this to be true for the IGHC genes. First, among the most remarkable findings from this cohort was the extraordinary number of novel IGHC alleles identified. Across the 9 core functional genes (and their newly curated paralogs) we identified a total of 221 coding alleles, 192 (86.8%) of which were not documented in IMGT. The IMGT database version we used for this paper (downloaded November 24, 2022) contained only 100 IGHC alleles. Thus, the collection of newly discovered alleles here represents a 192% increase. Importantly, many of these alleles carried changes in known functional residues, highlighting the potential for impacts on Fc function. Further to this point, the great extent of amino acid diversity described here opens opportunities to expand the identification of functional sites with novel functions.

A large fraction of the novel alleles identified here came from donors of non-European ancestries, stressing the ongoing need to generate improved genomic resources for the IG loci that are more representative of global populations. This is consistent with observations made in the V, D, and J genes for IGH, IGK, and IGL^13,37,41,42,54,59,60^. To contribute to this growing effort, data from this study and from our growing collection of long-read based assemblies will be made publicly available in VDJbase^59^ to enhance the representation of population diversity in existing IGHC germline allele databases. From VDJbase, users can browse and download annotated assemblies, allele sequences and usage statistics. Previously undocumented allele sequences are in the process of being submitted to the Nomenclature Review Committee of the International Union of Immunological Societies for ratification and naming.

Our long-read haplotype data also afforded us the ability to map these alleles to specific gene loci. To date, the majority of curated alleles have been identified and named, lacking knowledge and context of gene duplications, raising uncertainty in the gene assignments of existing alleles for genes with uncharacterized paralogs (Figure S16). A clear example is IGHG4 and related paralogs. Of the 13 alleles currently assigned to IGHG4 and the IGHG4 duplicate in IMGT, 10 of these alleles are assigned to IGHG4; however, whether these 10 alleles have been correctly assigned to their proper IGHG4 locus will need to be reassessed, deprecated and renamed when necessary. This is further complicated by the fact that, in addition to 3-copy IGHG4 haplotypes (which to date have not been discussed in the literature), we observed significant sequence variation among single-copy IGHG4 haplotypes including distinct deletion events harboring different IGHG4 paralogs. Thus, careful phylogenetic analysis will be required to account for this complex duplication structure and more fully delineate relationships between IGHG4 genes and alleles. Together, these observations illuminate significant flaws in our current nomenclature system.

An added strength of our comprehensive haplotype dataset is our sampling of donors from multiple global populations, allowing us to assess differences in IGHC haplotype variation between genetic ancestries. Previous IGHC gene sequencing studies have shown that allele frequencies vary significantly between human populations, and that coding variants exhibit high pairwise F_ST_ values between certain populations^17^. A genome-wide survey also found that a small 33 bp SV within IGHG4 was observed at a significantly higher frequency in Southeast Asian populations^20^. Here, we showed that these signatures extend to large SVs and SNVs in both coding and non-coding regions. Every common SV we assessed showed significant variation between populations. SV allele frequencies in AFR donors tended to deviate most from the other populations; for example, inversion SV genotypes were variable in all populations except AFR, in which only a single allele was observed in all samples. Similarly, AFR donors only carried single-copy IGHG4 haplotypes, whereas deletion haplotypes and 3-copy haplotypes were observed uniquely in EAS and EUR populations, respectively.

Additionally, a specific single-copy IGHG4 haplotype (cluster 4) was only observed in EAS and SAS, particularly enriched in EAS; this haplotype was associated with alleles most similar to IGHG4*08, and confirmed to represent haplotypes previously reported to be sites of local adaptation and introgression^20^ (Figure S17). Expanding population comparisons to locus-wide SNVs also demonstrated that both coding and non-coding variants contributed to population stratification. While the majority of IGHC intergenic and genic regions were associated with high F_ST_ estimates when comparing AFR to all other populations combined, smaller subregions of the locus surrounding only a subset of IGHC genes were associated with differentiating signatures for EAS and EUR as well. Interestingly, F_ST_ values were more modest in IGHD and IGHM regions, suggesting that these regions are potentially more homogeneous across genetic ancestries.

Future work should seek to understand how this extensive genetic diversity influences the composition of the expressed Ab repertoire and contributes to variation in Ab effector function. Allelic diversity has been demonstrated to influence functional outcomes by altering FcR and IG interactions, thereby affecting Ab half-life^25^. Additionally, SVs have been linked to differential ADCC and enhanced complement activation, including IGHG3 hinge copy number^61,62^. IGHG4, which we have shown boasts extensive copy number variation differences across different ancestries, is generally associated with low inflammatory activity and weak complement activation. However, coding variants of IGHG4 have been reported to be associated with elevated expression levels, as well as variation in Fab arm exchange and patterns of glycosylation^53,63^. In addition to coding variation, it will also be worth exploring whether non-coding variants influence biases in CSR frequencies, leading to shifts in the distribution of particular IGHC genes and linked IGHV, D, and J transcripts. Our recent studies of near-full length IgM and IgG transcripts^13,48^ have revealed that multi-copy IGHC genes (such as IGHG4 and IGHA1 and their close paralogs) are detected in the Ab repertoire. Using our recently developed long-read based approaches will be important for identifying interindividual variation in the Ab repertoire, accounting for full BCR transcript variation to identify potential disease-associated signatures.

In conclusion, our population-level survey has revealed that the human IGHC locus harbors extensive genetic diversity, including variants that differentiate human populations, with the potential of contributing to Ab function and disease.

### Limitations of the Study

Together our data provide evidence that IGHC diversity has been vastly underestimated and is deserving of more focused study. Specifically, the population signatures we have observed in this study may point to historical demographic events, geographic isolation, co-evolution with pathogens, and/or variation in selective pressures. However, it is important that we reiterate the relatively small subpopulation sample sizes collected here. This limitation demands further analysis in larger cohorts to both validate our observations and investigate these scenarios.

Continued efforts to characterize haplotype variation at scale will be needed to delineate functional variants of biomedical importance. We show here that the use of long-read sequencing offers a tractable solution to this challenge. However, it will be important to increase the capacity of existing molecular and analysis tools to allow for more automated assessments of long-read based assemblies, with respect to their contiguity and phasing accuracy, as well as ensuring precision in SV and SNV discovery and genotyping. Finally, the discovered haplotype complexity also demands greater attention to the development of pangenome reference resources that can better represent haplotype variation^64–67^. Moving forward, these will allow for the more ordered cataloguing of IGHC genetic variants of all types, as well as the curation of genes and alleles with new and improved nomenclatures^68^ that more aptly reflect the population and evolutionary history of these loci.

## Resource Availability

### Lead contact

Further information and requests for resources should be directed to and will be fulfilled by the lead contact, Corey T. Watson.

### Materials availability

This study did not generate new unique reagents.

### Data and code availability

All raw sequencing data in BAM format have been deposited in SRA and are publicly available as of the date of publication. Allele sequences, corresponding assemblies and alignment files will be made available at https://vdjbase.org/ with sample metadata. Raw data is deposited in SRA under BioProject PRJNA555323, accessions listed under the key resources table. All original codes have been deposited in Zenodo (https://doi.org/10.5281/zenodo.15802507).

## Acknowledgments

We sincerely thank Eric Rouchka and Juw Won Park for their valuable insights and feedback on the computational analysis. We also appreciate the contributions of Easton Ford, Kamille Rasche and Chandrima Bharadwaj for their assistance with manuscript preparation, and a special thanks to Andy Lauer and the staff of the UofL Sequencing Technology Center for assistance in the sequencing.

## Author contributions

U.J., O.L.R., M.L.S., and C.T.W. contributed to the conception of the work. M.L.S. directed wet lab/bench and sequencing experiments. K.S., S.S. prepared sequencing libraries. U.J., O.L.R., W.L., E.E., and Z.V. developed the computational pipeline to analyze the sequencing data. U.J., O.L.R., W.L., E.E., W.S.G., G.Y., C.T.W. interpreted results. C.T.W., M.L.S. supervised the experiments, analysis and data interpretation. C.T.W., M.L.S. and E.E.E. acquired funding. U.J. and C.T.W. drafted the manuscript and all authors contributed to revising the manuscript.

## Declaration of interests

C.T.W., M.L.S., and W.L. are founders and shareholders of Clareo Biosciences, Inc. and serve on its Executive Board. M.L.S. and C.T.W. are listed inventors of patent filing PCT/US2024/044692. E.E.E. is a scientific advisory board (SAB) member of Variant Bio, Inc. R.S. is co-founder and CEO of Panacent Bio, Inc. None of the other authors declare any competing interests.

## Funding

This work was supported by an award made to C.T.W. and M.L.S. by the NIAID (R24 AI138963) and in part to E.E.E. by the US NIH (R01HG010169). E.E.E. is an investigator of the Howard Hughes Medical Institute.

## Star Methods

### Key resources table

#### Chemicals, peptides, and recombinant proteins

|  |  |  |
| --- | --- | --- |
| G-tubes | Covaris | Cat#520104 |
| Buffer EB | Qiagen | Cat#19086 |
| AMPure PB Beads | Pacific Biosciences | Cat#100-265-900 |
| Barcode adapters, DNA Plate Duplex Oligo, 50 µM | IDT | IDT Plate #13748866 |
| KAPA Pure Beads, 60 mL | KAPA Roche | Cat#07983298001 |
| Universal Oligos, 100 µM | IDT | Ref#288647790 |
| KAPA Probes Resuspension Buffer | KAPA Roche | Cat#09075879001 |
| 2.0 mL Deep Well Plate | Genesee Scientific | Cat#27-413 |
| LB Agar Powder | VWR | Cat#89233-940 |
| Chloramphenicol | VWR | Cat#45000-618 |
| DL-Arabinose | VWR | Cat#IC10070010 |
| Buffer P1 | Qiagen | Cat#19051 |
| Buffer P2 | Qiagen | Cat#19052 |
| Buffer P3 | Qiagen | Cat#19053 |
| Isopropanol | Fisher Scientific | Cat#AC423830010 |
| TE Buffer | Thermo Fisher Scientific | Cat#12090015 |

#### Critical commercial assays

|  |  |  |
| --- | --- | --- |
| Qubit 1x dsDNA BR Assay kit | Thermo Fisher | Cat#Q33266 |
| 0.75% Agarose cassettes, Dye Free, Low Range 10/pk S1 | Sage Science | Cat#BLF7510 |
| KAPA Hyper prep kit, 96 rxn | KAPA Roche | Cat#07962363001 |
| PrimeSTAR® GXL DNA Polymerase, 1000U | Takara Bio | Cat#R050B |
| DNF-467 Genomic DNA 50 kb kit | Agilent | Cat#DNF-467-0500 |
| KAPA HyperCapture Reagent kit, 96 rxn | KAPA Roche | Cat#09075828001 |
| KAPA HyperCapture Bead kit, 24 rxn | KAPA Roche | Cat#09075780001 |
| SMRTBell Express Template Prep Kit 2.0 | Pacific Biosciences | Cat#100-938-900 |
| SMRTBell Enzyme Cleanup Kit 1.0 | Pacific Biosciences | Cat#101-746-400 |

#### Deposited data

|  |  |  |
| --- | --- | --- |
| Raw sequencing data | BioProject PRJNA555323 (This paper) | SRA accessions: SRR32268925 - SRR32269029, SRR32483169-SRR32483171 |
| Assembly, annotation | Genbank (This paper) | PV296382-PV297760, PV329667- PV329688, PV359020-PV359166 |

#### Databases and software

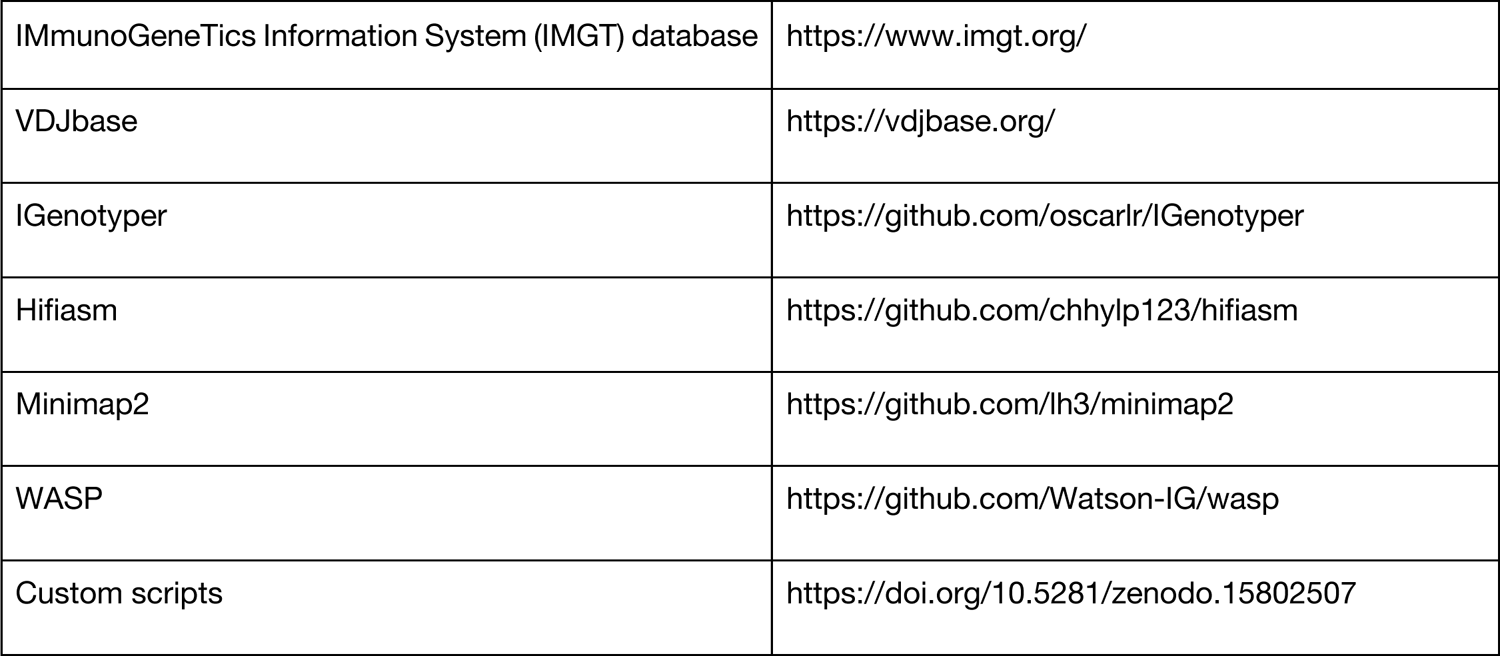

### Method Details Sample information

Samples used in this study were previously collected as a part of the 1000 Genomes Project (1KGP)^69^. Genomic DNA was procured from the Coriell Institute for Medical Research (https://www.coriell.org/; Camden, NJ). Fosmid libraries were originally generated from lymphoblastoid cell lines (LCL)^70^. Sample, population, sub-population, sex, and relatedness information are reported in Table S10. While EBV (Epstein-Barr virus) transformed LCLs are valuable for studying human genetic diversity, the IGHC loci were carefully curated to account for class switch recombination, ensuring DNA integrity and haplotype preservation.

### Constructing a reference assembly haplotype for the human IGHC locus

To build a reference assembly for the mapping and characterization of genetic variants in samples sequenced in our study, we utilized the IGHC locus from the T2T CHM13 v2.0 build (chr14:99730306-100132055) as our IGHC reference for all analyses conducted below. We chose to use this assembly rather than GRCh38/hg38 because it harbored a ∼19.5 Kbp insertion (Figure S18), including a duplication of the gene IGHG4.

### Fosmid isolation, sequencing and assembly

Fosmid clones were selected based on clone-end sequence mapping to the IGHC locus (GRCh38; chr14:105,510-105,860,000). For each individual, five non-overlapping pools of fosmids were selected (average n=16 per pool). Fosmids were grown in 96-well plates and DNA was extracted following previously published methods^70^. On a per fosmid clone basis, DNA was sheared using g-tubes (Covaris, Woburn, MA, USA) to ∼15 Kbp, followed by size selection of >10 Kbp fragments using the BluePippin system (Sage Science, Beverly, MA, USA). Following shearing and size-selection, DNA from non-overlapping fosmids was pooled in equimolar ratios, with the number of samples per pool dependent on the SMRT sequencing platform used. Pooled DNA was then directly prepared for sequencing using SMRTbell Template Preparation Kit 1.0 (Pacific Biosciences, Menlo Park, CA, USA) following the manufacturer’s standard protocol. SMRTbells were sequenced on the RSII sequencing system (Pacific Biosciences) using P6/C4 chemistry and 6 hour movies. Fosmid-level metadata are provided in Table S11.

For each fosmid pool, sequencing data from individual fosmids were used to generate fosmid-specific assemblies by first removing the fosmid backbone sequence from the reads and mapping them to the GRCh38 reference. Then, for each fosmid, only reads that mapped within the fosmid end sequence coordinates were de novo assembled using Canu^47,70^. Clones with ends that did not map well to the reference genome (∼40 Kbp chunks) were marked discordant; those were barcoded and sequenced. These fosmids were assembled *de novo* without mapping.

### Targeted capture sequencing and assembly

For targeted long-read sequencing of the IGHC locus, we designed a tailored HyperCap DNA probe panel (Roche). This panel included target sequences from the GRCh38 IGHC locus (chr14:105,510-105,860,000). Genomic DNA (gDNA) samples were prepared and sequenced according to previously published methods^36,38,41,42^. Briefly, gDNA was sheared to approximately 14.4 Kbp (median) using g-tubes (Covaris, Woburn, MA, USA) and size selected to remove fragments less than 3-5 Kbp, using either bead-based size selection or with the Blue Pippin system (Sage Science, Beverly, MA, USA). Fragmented gDNA was ligated to universal adapters and amplified. Amplified individual samples were pooled in groups of 6 prior to target-enrichment using the custom HyperCap DNA probes. After probe capture, targeted fragments were subjected to a second round of amplification and processed for SMRT sequencing using the SMRTbell Template Preparation Kit 2.0 as per the manufacturer’s protocol. The resulting SMRTbell libraries were multiplexed across pools as 2-plex (12 samples) and sequenced using one SMRT cell 8M on the Sequel IIe system with 2.0 chemistry and 30-hour movies.

Following data acquisition, HiFi (high-fidelity; > Q30 or 99.9%) reads were generated and aligned to the IGHC reference for coverage analysis to first assess sequencing depth over the IGHC region; non-specific reads mapping to non-IGHC regions were discarded; the mean per-base read coverage for each sample is provided in Table S10. HiFi reads were used to generate phased *de novo* assemblies using Hifiasm with default parameters^46^. Assemblies were aligned to the IGHC reference using minimap2 (version 2.26)^71^ with the ‘-x asm20’ allowing maximum divergence in alignments and using custom gap penalty adjustments to accommodate better alignments over structural variants. HiFi reads were also used to generate orthogonal assemblies using the IGenotyper^38^ pipeline with default parameters to generate reference-guided locally phased assemblies. The Hifiasm-based assemblies, which offer greater contiguity, were aligned to the reference and utilized for SV detection. Subsequent analysis involved SV annotation and allele curation across SV regions using haplotype-specific Hifiasm contigs. Assemblies generated by IGenotyper were guided by the reference genome and phased using contiguous read support in conjunction with the presence of heterozygous single SNPs. Phasing blocks were defined as the reference span between the two farthest SNPs connected by contiguous read support, and these blocks were utilized to generate the most accurate haplotype-specific contigs. These contigs were then aligned back to the reference genome to obtain contig alignment over the IGHC locus, excluding the IGHG4 SV region. The alignments were then examined for areas with missing contigs using the bedtools genomecov command, confirming the presence of two heterozygous contigs (hap=1, hap=2) or one homozygous contig (hap=0) in every alignment block. To fill in these alignments, curated missing region boundaries were supplemented using *de novo* assemblies generated by Canu^47^. These assemblies were created based on available HiFi reads aligned specifically to the missing region boundary, resulting in the production of missing haplotypes or alternate haplotypes to complete the final assemblies. These contigs were also aligned against the IGHC reference using minimap2 (version 2.26) with the ‘-x asm20’ for allowing maximum divergence in the alignment and using custom gap penalty adjustments as mentioned before.

### Construction of hybrid assemblies

Fosmid contigs were aligned to the IGHC reference for each individual and phased into haplotype groups using overlapping SNP profiles detected through manual inspection of contigs in the Integrative Genomics viewer (IGV)^72,73^. Haplotype-specific fosmid contigs were merged to form larger contigs using a custom python script, requiring a minimum 2 Kbp overlap, with a maximum 5 bp mismatches. Gaps in the resultant merged assemblies were filled using assemblies generated from long-read targeted sequencing from the same donors (Table S12). To validate bases in hybrid assemblies, HiFi reads from IGHC locus-wide targeted capture sequencing from the same donor were realigned to each diploid assembly. Base discrepancies between reads and assembly bases were assessed using the WhatsHap^74^ find_snv_candidates command to detect SNPs potentially representing erroneous bases in these diploid reference assemblies.

### Construction of targeted capture phased assemblies in trio probands

We selected 4 human trios for which we generated targeted long-read sequencing, and for which the proband dataset had a mean 30X read depth over IGHC locus bases. The parental HiFi reads were processed using Yak to generate a count of k-mers present in each parent, utilizing the parameters “-k31 -b37”. Subsequently, the proband was assembled using Hifiasm in trio binning mode^46^, employing the parental k-mer count. Additionally, the proband HiFi reads were assembled using IGenotyper^38^. The proband contigs obtained from the Hifiasm and IGenotyper pipelines were inspected in IGV^72,73^ alongside their respective parental Hifiasm contigs to ensure that phased contigs in the proband were supported by those generated in the parents. Any phase switch errors in Hifiasm contigs were corrected using IGenotyper contigs to produce final gapless contigs for each haplotype of the proband. Proband contigs derived from trio-based analyses underwent orthogonal validation, employing a method similar to that used for hybrid assemblies. This involved realigning capture reads to their personalized diploid references and assessing the presence of erroneous bases within the diploid reference using SNP calls obtained through the WhatsHap find_snv_candidates command.

### Detection and genotyping of IGHC single nucleotide variants in phased hybrid and proband assemblies

Samples with hybrid assemblies and trio-based proband assemblies were processed using ‘bcftools mpileup’^75,76^ command with ‘-Ou -f -B’ to call variants and ‘-r’ was used to specify the region of targeted capture alignment, for the HiFi reads based on capture probes, then ‘bcftools call’ command was used with ‘-m -Ov-V indels’ to get all bases reported alongside variants, skipping the indels. The ‘bcftools view’ command was used with “ -i ‘QUAL >= 0’” to filter positions with zero quality scores, then the ‘bedtools intersect’^77^ command was used with ‘-b masking_regions.bed’, which consists of common IGHC SV coordinates to skip any entries in the vcf file over SV regions. These vcf files were used as ground truth to assess targeted capture-only diploid assemblies processed through our custom IGHC genotyping pipeline.

To analyze population statistics, variant data from targeted capture-based assemblies with a MAF > 5% were utilized. The R package SNPrelate^78^ was utilized to perform PCA, enabling the assessment of population relatedness using identity-by-descent (IBD) measures. Pairwise F_ST_ analysis was conducted between a specific population group and the rest of the individuals using the same variant dataset to evaluate genetic differentiation. Weir and Cockerham’s F ^57^ values were calculated for all IGHC SNPs having a minor allele frequency >5% using VCFtools v0.1.16^79^.

### Benchmarking the accuracy of high-throughput IGHC targeted long-read sequencing in diploid samples

To create test SNV genotype call sets for benchmarking, targeted capture assemblies for the same individuals (fosmid donors and trio probands, as described above) were processed through both the Hifiasm and IGenotyper pipelines. The contigs generated by each computational pipeline were aligned to the IGHC reference to produce assembly alignments (bam files). These bam files were subsequently processed to call SNVs, after masking common SV regions, such as the IGHG4 and IGHG2-IGHA1 inversion regions, to avoid confounding the results. For the highly contiguous Hifiasm assemblies, SNVs were curated using ‘bcftools mpileup’^76^. In the case of IGenotyper assemblies, variant calls were first made directly from the assembly contigs aligned against the reference genome using ‘bcftools mpileup’, followed by filtering of this call set to only include SNVs supported by HiFi read pileups. To do this, HiFi reads were processed using two approaches: SNP calling via ‘bcftools mpileup’^76^ and ‘whatshap find_snv_candidates’ with the ‘--minrel 0.05’ option^74^, which includes all variant positions while masking SV regions. The HiFi-based vcf file was constructed by combining SNV positions identified by whatshap and entries from bcftools mpileup, with overlapping variants resolved by prioritizing whatshap calls to create the final HiFi-reads-based vcf. The final HiFi-based vcf and capture-assembly-based vcf were then intersected to produce the most accurate set of variant calls for each individual. Finally, genotype calls from the capture-only assemblies were compared against ground truth sets to evaluate the performance of the high-throughput pipeline.

### Benchmarking assembly approach using variable length synthetic reads from vetted assemblies

To evaluate the accuracy of assemblies using different input read lengths, we simulated reads from two benchmark diploid assemblies and the IGHC reference haplotype. In each iteration, a custom Bash script simulated random reads of 350 bp, 3 kb, 6 kb, and 10 kb with overlapping boundaries, limited to 15× coverage per haplotype. For diploid samples, reads from both haplotypes were merged and assembled using Hifiasm^46^. Assemblies generated using synthetic reads were then compared directly to source assemblies. Accuracy was assessed by calculating average error rates in 10 kb windows across read length bins to identify the optimal read length for resolving target regions.

### Identification and genotyping of structural variants

For structural variants in the IGHC region, assembly bam files were manually inspected in IGV. In this study, SV genotyping was mainly focused on the most commonly found SVs in the IGHC region resolved by the assembly process. Alignment files from IGenotyper and Hifiasm assemblies were manually inspected in IGV to identify large deletions (IGHG4D), inversions (between IGHG2 and IGHA1), and hinge deletions (IGHG3), which were then genotyped accordingly. Additional large SVs identified in only a small number of individuals were identified and curated through manual assessment of the assembly contigs (Figure S6). From our targeted capture assemblies we initially observed the presence of a third contig over the IGHG2-IGHA1 region, suggestive of a duplicated sequence. To be certain with the observation we also assessed the publicly available long-read whole genome sequencing (WGS) data (Pacific Biosciences) of these individuals from the Human Pangenome Reference Consortium (HPRC) and confirmed similar instances from soft clipped and supplementary alignments. From the alignment on the current reference assembly over the IGHE-pseudogene (IGHEP1) region, manual inspection of the plausible duplication sequence for putative INDELs (2-3 Kbp) and review of the softclipped unaligned portion using the blat toolkit in the UCSC genome browser revealed alignment hits over IGHE and/or IGHG4, hinting at the large segmental duplication (segdup) structure previously described by^15^. We further used a 150k-sized k-mer based read simulation from the WGS contigs to confirm the plausible breakpoint using the GRCh38 IGHC reference as the guiding frame (Figure S6). Once confirmed we separated the contigs and used the haplotype-specific contigs from both WGS and targeted capture to merge them based on terminal overlap and sequence identity, producing the complete contiguous haplotype for the duplication and alternate haplotype, respectively. Further, we used these haplotypes as a personalized reference for realignment of the HiFi reads for orthogonal validation of the SV.

To generate genotypes for the complex IGHG4 region, we extracted sequences from each haplotype-resolved Hifiasm assembly spanning the IGHG4 region with a flanking sequence length of +/- 5 Kbp. This was followed by a length binning approach based on tandem repeat size to ensure comparison among similar copy number variation (CNV) haplotypes, and then multiple sequence alignment was performed for all haplotypes in our cohort. From this alignment, a pairwise distance-based matrix was formulated, followed by hierarchical clustering to identify IGHG4 region spanning haplotype clusters. The distance threshold (0.1) determined the number of clusters to be assigned to the haplotypes. Individuals were then genotyped based on the identified haplotype clusters present in each donor.

### Identification of IGHC alleles

Allele sequences for the IGHC genes were extracted from haplotype grouped IGenotyper and Hifiasm assembly alignments using a custom script (https://github.com/Watson-IG/wasp) to generate an annotation file for all the genes in each sample. IGHC alleles were assigned IMGT allele names for identical sequence matches. Alleles with <100% identity alignments were assigned to the closest IMGT allele, followed by designation of polymorphic positions following the VDJbase nomenclature schema (https://vdjbase.org/). For IGHC genes that were determined to be outside of an SV in a given sample, allele sequences were extracted using the IGHC_gene_coords.bed based on the alignment of a given assembly to the IGHC reference, requiring that a given extracted allele had to include all exons within a single contig (Table S13, S14). The reference database used for the assignment of “known” and “novel” alleles was downloaded from IMGT^19^ in November 2022.

Support from HiFi reads for curated alleles from each assembly contig was assessed using a custom read-support script (https://github.com/Watson-IG/wasp/tree/master/annotation/read-support). Specifically, donor-level HiFi reads were realigned to the respective contig assemblies generated for that donor. For each allele, the number of fully spanning HiFi reads was enumerated, counting the number of HiFi reads with 100% base agreement with the curated allele across all exons in the personalized alignment. In addition, read support was calculated for each base position of the curated allele from all reads aligned to the personalized reference, regardless of whether that read fully spanned the gene. The percentage of reads supporting the base at that position in the assembly was recorded (using samtools mpileup;^76^). We set our read support threshold to require that at least 10 reads mapped to all exonic positions and that at least 80% of those reads supported the base represented by the assembly and curated allele.

### Identifying amino acid changes in functional residues

To evaluate the potential functional impact of curated IGHC alleles, we compiled a list of functionally important residues from literature sources reporting altered Fc-FcγR and Fc-FcRn binding^7,25^, changes in complement activation^53^, ADCC activity^26,80^, and modifications affecting antibody half-life^81–83^. These residues were cross-referenced against gene-specific multiple sequence alignments (MSAs) generated using Clustal Omega^84^ for each IGHG and IGHA isotype. Allelic variants were mapped to these annotated positions to identify amino acid substitutions with known or potential functional relevance.

To further assess structural implications, variants at mapped functional sites were modeled using homology-based structural modeling via SWISS-MODEL^85,86^, leveraging relevant antibody–receptor or antibody–ligand complexes from the Protein Data Bank (PDB)^87,88^. Structural models were analyzed for changes in bonding interactions, bond lengths, and physicochemical properties to infer possible effects on molecular function.

## Supplementary information

Document S1. Figures S1–S18 (supplementary_figures.pdf)

Document S2. Tables S1–S17 (supplementary_tables_CG_v3.xlsx)

Tables S1-4. Construction of primary vetted assemblies and benchmarking on computational assembly approaches, related to Figure 3 and STAR Methods

Tables S5-6. Evaluation of concordance between long-read variant calls and reference databases such as 1KGP and dbSNP, related to Figure 4

Tables S7-8. Potential functional impacts of IGHC variants in expressed alleles, related to STAR Methods and Figure S13

Table S9. IGHG4 deletion haplotype alleles and corresponding clusters, related to Figure 6 and S14 Tables S10-14. Distribution of ethnicities and alleles across individual samples, related to STAR Methods

Tables S15-17. Variations among frequently observed alleles compared to the consensus allele within the cohort (n=105), related to Figure 5 and S11, S12

## Notes

### Competing Interest Statement

The authors have declared no competing interest.

### Summary of Updates

This version of the manuscript has been revised to update "Title" and overall word limit (except Methods) as per journal's suggested limits (145 characters and 8200 words respectively)

