## Supplementary Figures for "The human IG heavy chain constant gene locus is enriched for large structural variants and coding polymorphisms that vary among human populations"

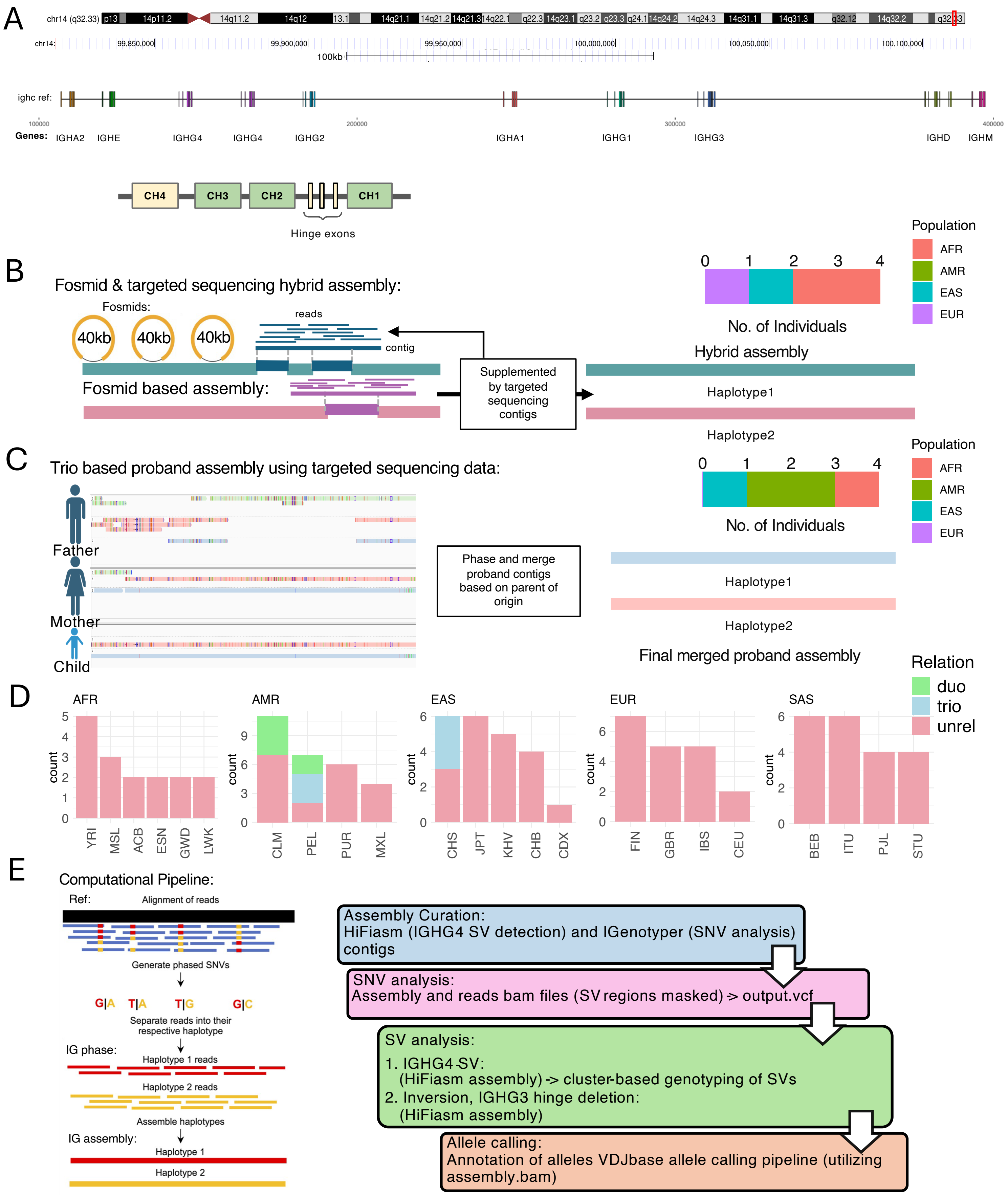


**Figure S1. A comprehensive overview of methods and samples used for analyzing IGHC genomic diversity, related to STAR Methods**

**(A)** schematic map of the IGHC locus and gene structure. **(B-C)** Schematic representation of the selected ground truth assemblies - fosmid, long-read targeted capture data was used to produce highly accurate hybrid assemblies; targeted long read sequence from both parents and proband (trio) was used to produce the second set of highly vetted assemblies. **(D)** A breakdown of relation and ancestral information for the 105 individuals used in the study. **(E)** The schematic representation of high-throughput computational pipeline used for analyzing IGHC locus.

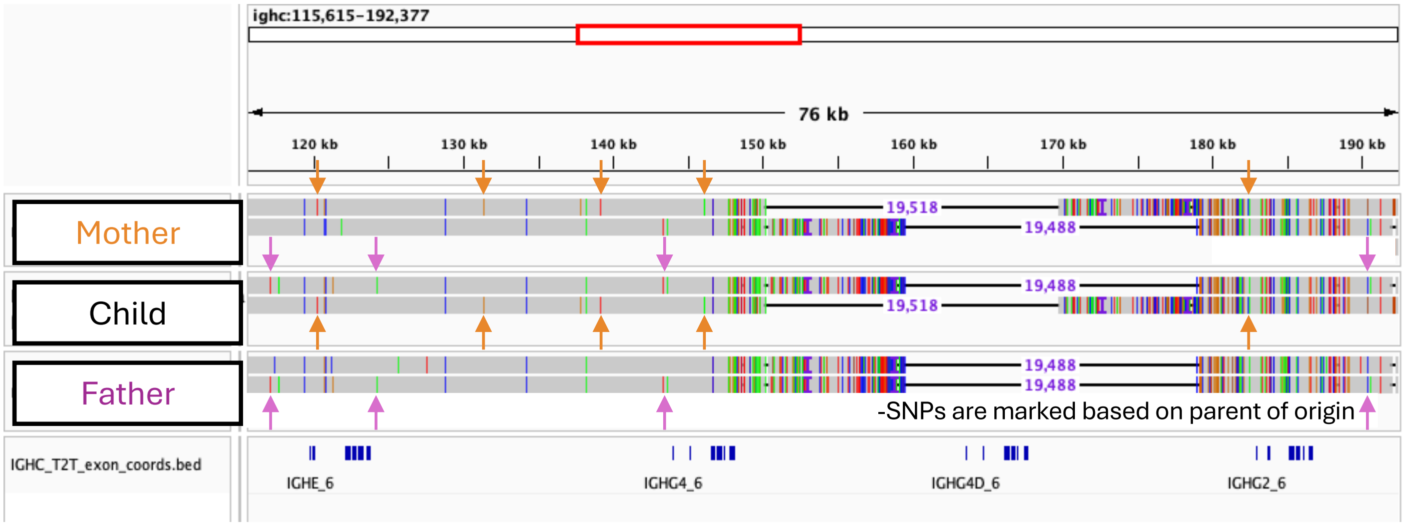


Observing child assembly alongside parental assemblies :

- Orange arrow marked SNPs are maternal haplotype guide within the proband contig
- Purple arrow marked SNPs are paternal haplotype guide within the proband contig
- Parallel observation of paternal and maternal contigs for ensuring accurately phased proband contigs

**Figure S2. A schematic representation for manually ensuring correct phasing in trio-based proband assemblies, related to STAR Methods**

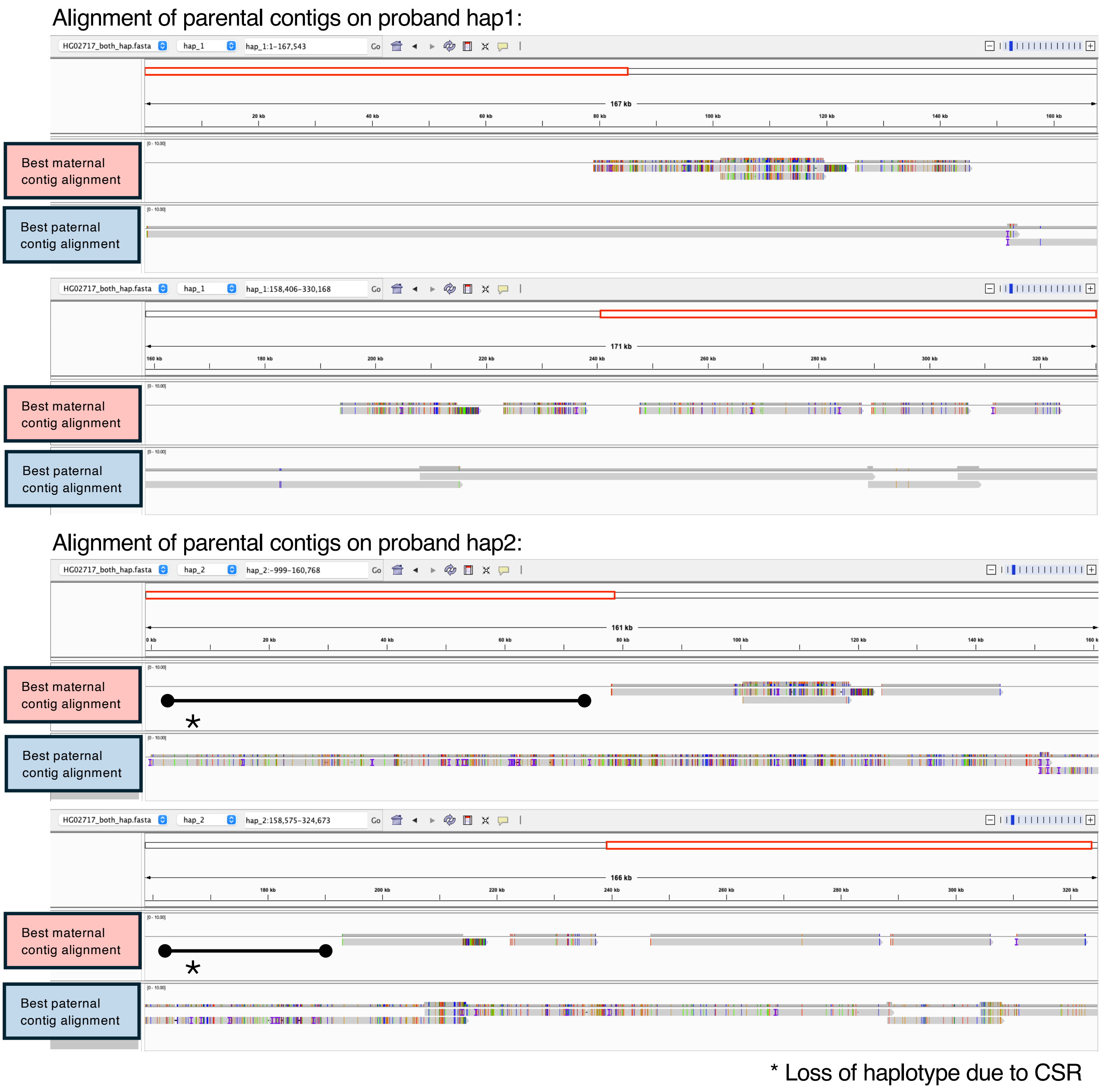


**Figure S3. Representation of correct phasing in scenarios harboring loss of haplotype, related to STAR Methods**

Illustrating here is a Haplotype loss as observed in the mother's assembly (HG02716); likely due to a class-switch event. However, complete paternal assembly (HG02715) and partial maternal assembly data proved adequate for reconstructing the proband assembly (HG02717).

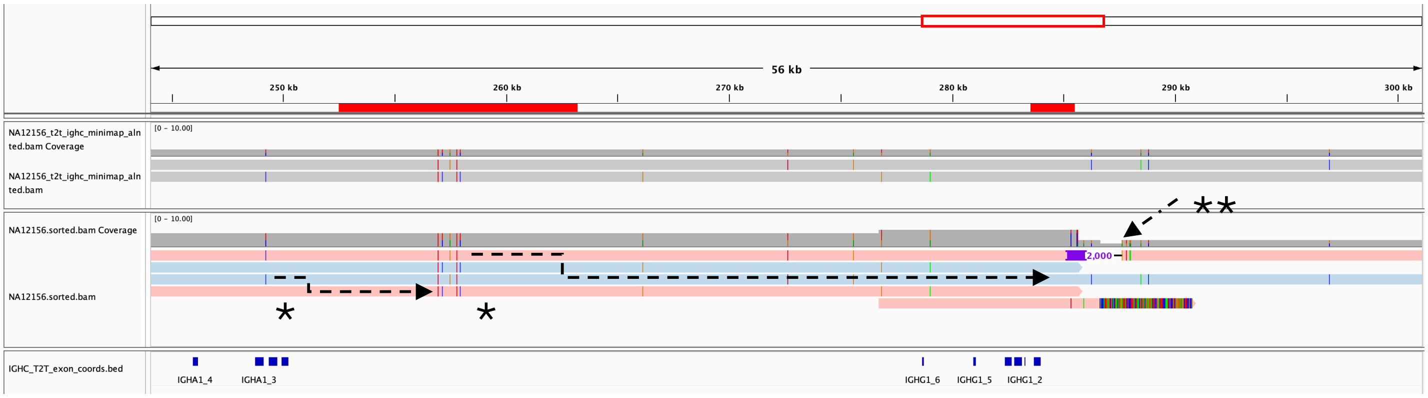


* Phase switches

** assembly error due to class switch recombination

**Figure S4. An example representation of phase switch errors and mis-assembly caused by class-switch recombination, related to Figure 3 and STAR Methods**

The dotted arrows (marked with *) connecting haplotypes indicate the correct phasing over the shown haplotypes. The contiguous assembly contig over the region (marked with **), with an exact haplotype representing contig with right soft-clipping, points to the presence of a class-switch event (DNA rearrangement placing VDJ sequences in position) after IGHG1, which results in mis-assembly in the primary assembly contigs.

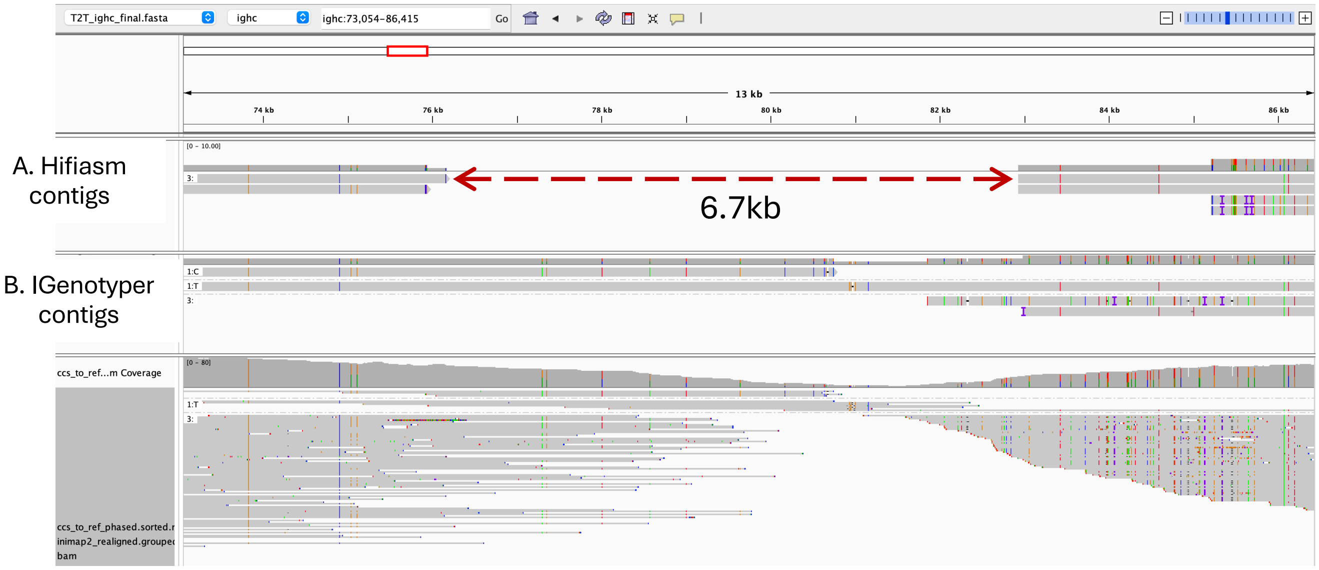


**Figure S5. An example of a comparison for assembly contig generation between IGenotyper and Hifiasm, related to Figure 3 and STAR Methods**
IGenotyper extends/produces contig over lower depth, whereas Hifiasm skips the extension of the contig at such regions.

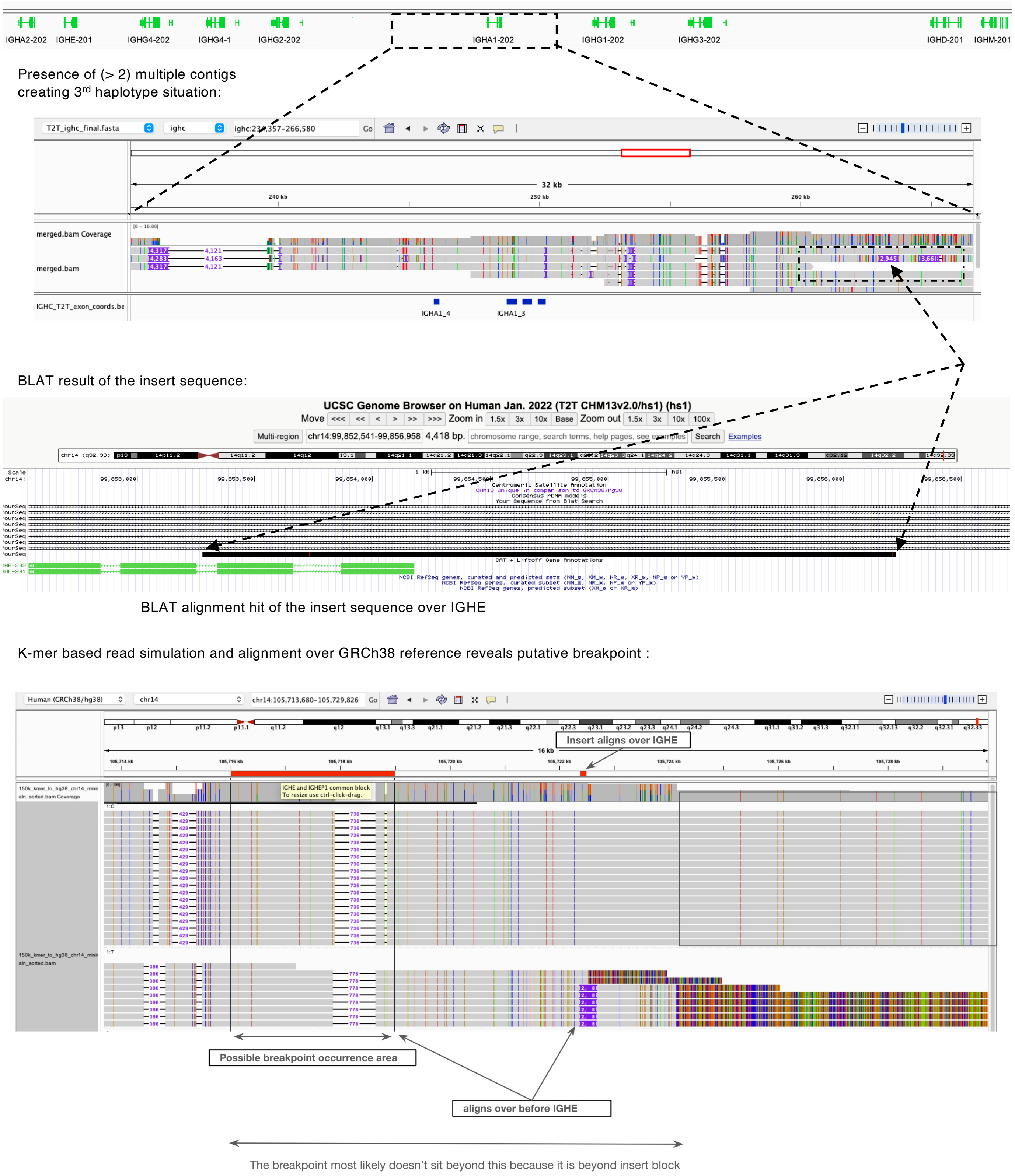


**Figure S6. Identification of break points in large segmental duplication block observed in the Hifiasm de-novo assembly contigs, related to Figure 4 and STAR Methods**

The presence of a large segmental duplication was confirmed through the identification of a third haplotype within the alignment. Subsequently, the mapping locations of the inserted sequences (~2–3 kb) were determined using the UCSC Genome Browser BLAT toolkit. The BLAT results revealed alignment over IGHE rather than the pseudogene (EP), providing initial insights into the potential breakpoints. Furthermore, a read simulation approach was employed using the resolved duplicated haplotype to generate 150 kb-sized k-mers as reads. These simulated reads were then aligned to the reference genome to gain a clearer visual representation of the putative breakpoint.

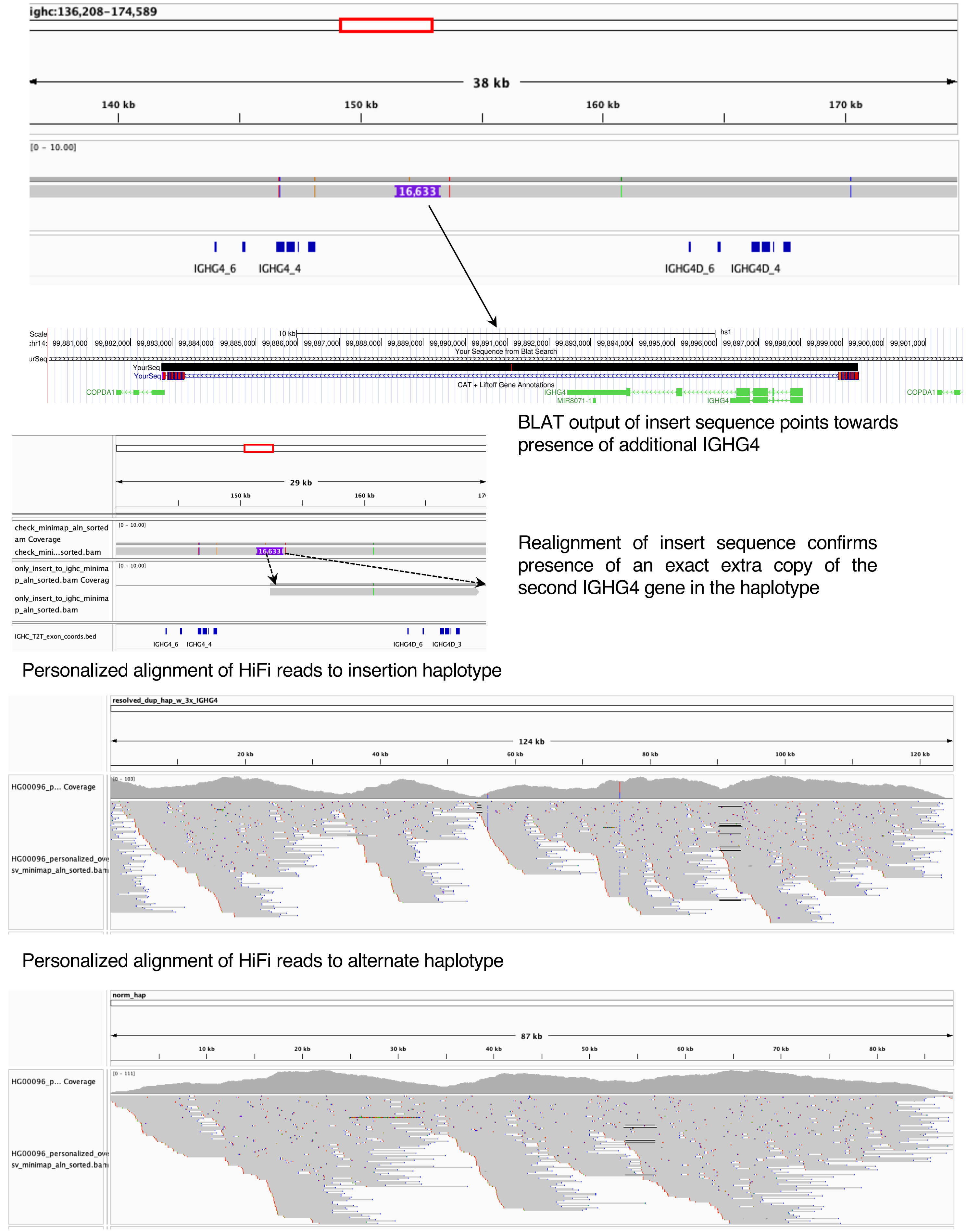


**Figure S7. Mapping of 3CNV IGHG4 haplotype over IGHC reference from T2TCHM13v2.0, related to Figure 4 and STAR Methods**

The EUR haplotype contains two copies of IGHG4, identical to the reference, along with a ~16kb insert between the genes. Subsequent alignment of the insert sequence reveals the presence of a third copy.

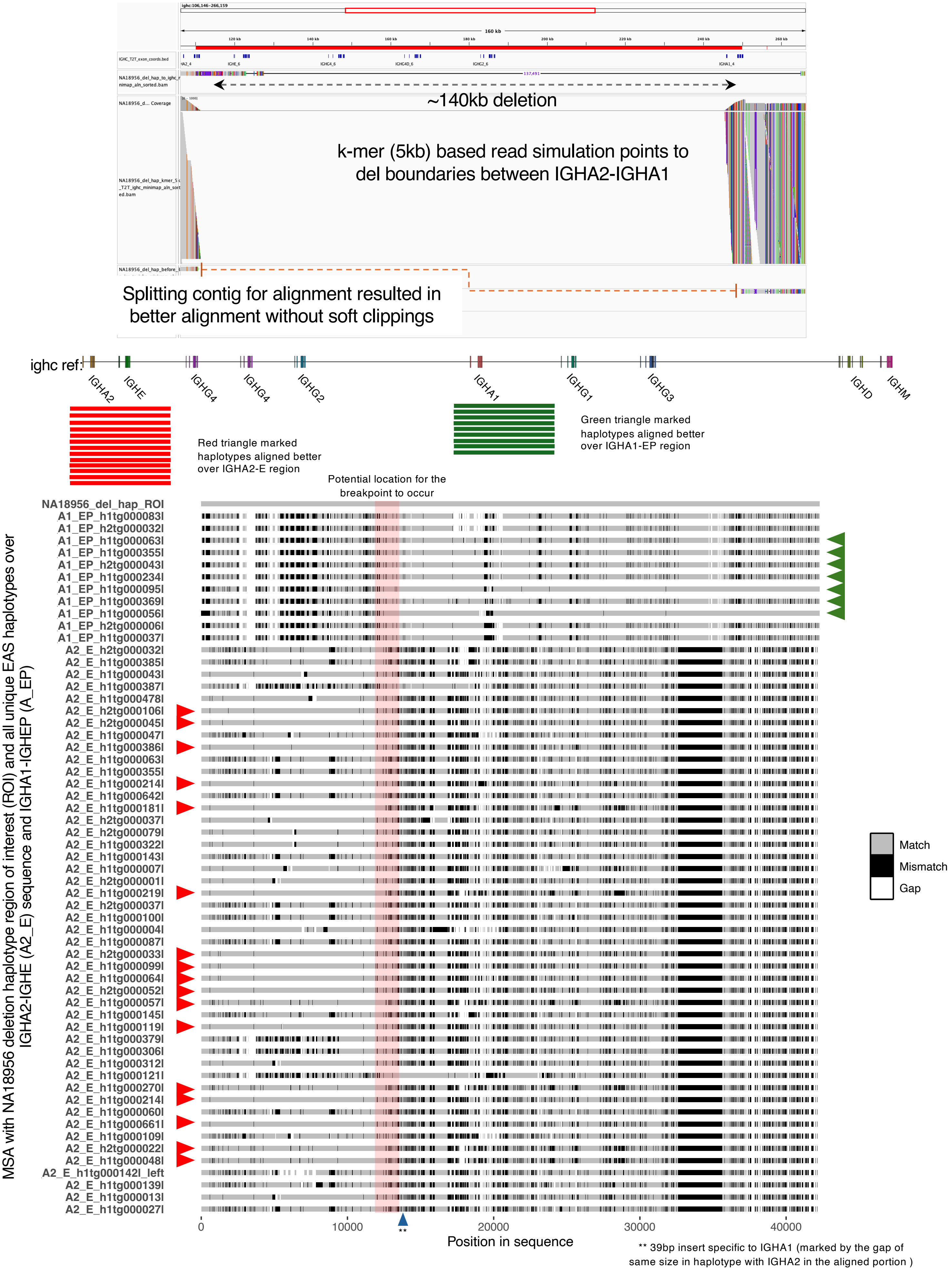


**Figure S8. Schematic for potential breakpoint estimation in the multigene deletion haplotype (NA18956), related to Figure 4 and STAR Methods**

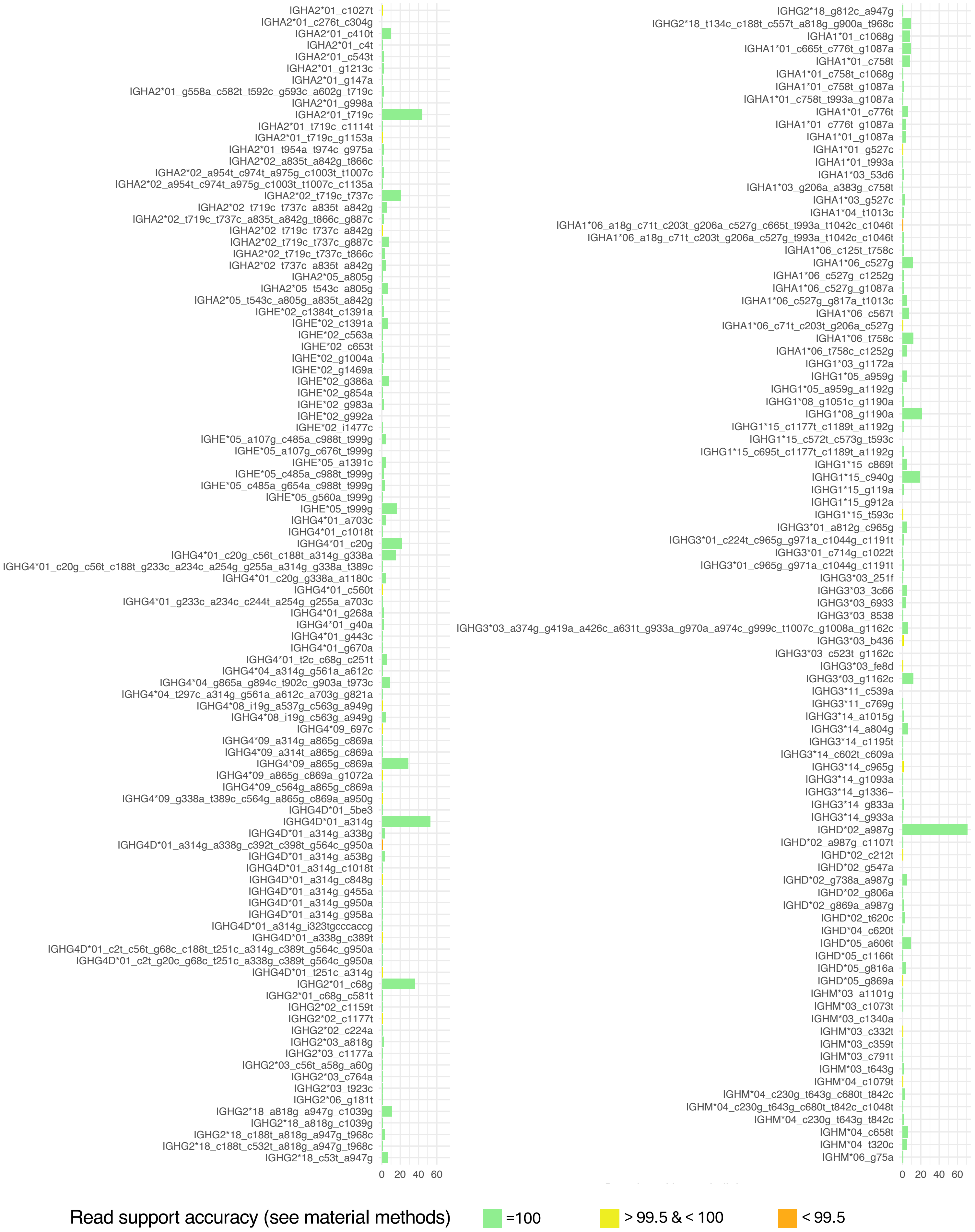


**Figure S9. Bar plot for frequency distribution of novel alleles observed in this study, related to Figure 5**
Here "X" axis indicates no. of individuals observed with the novel allele and the color of the horizontal bar indicates read based accuracy support for the allele

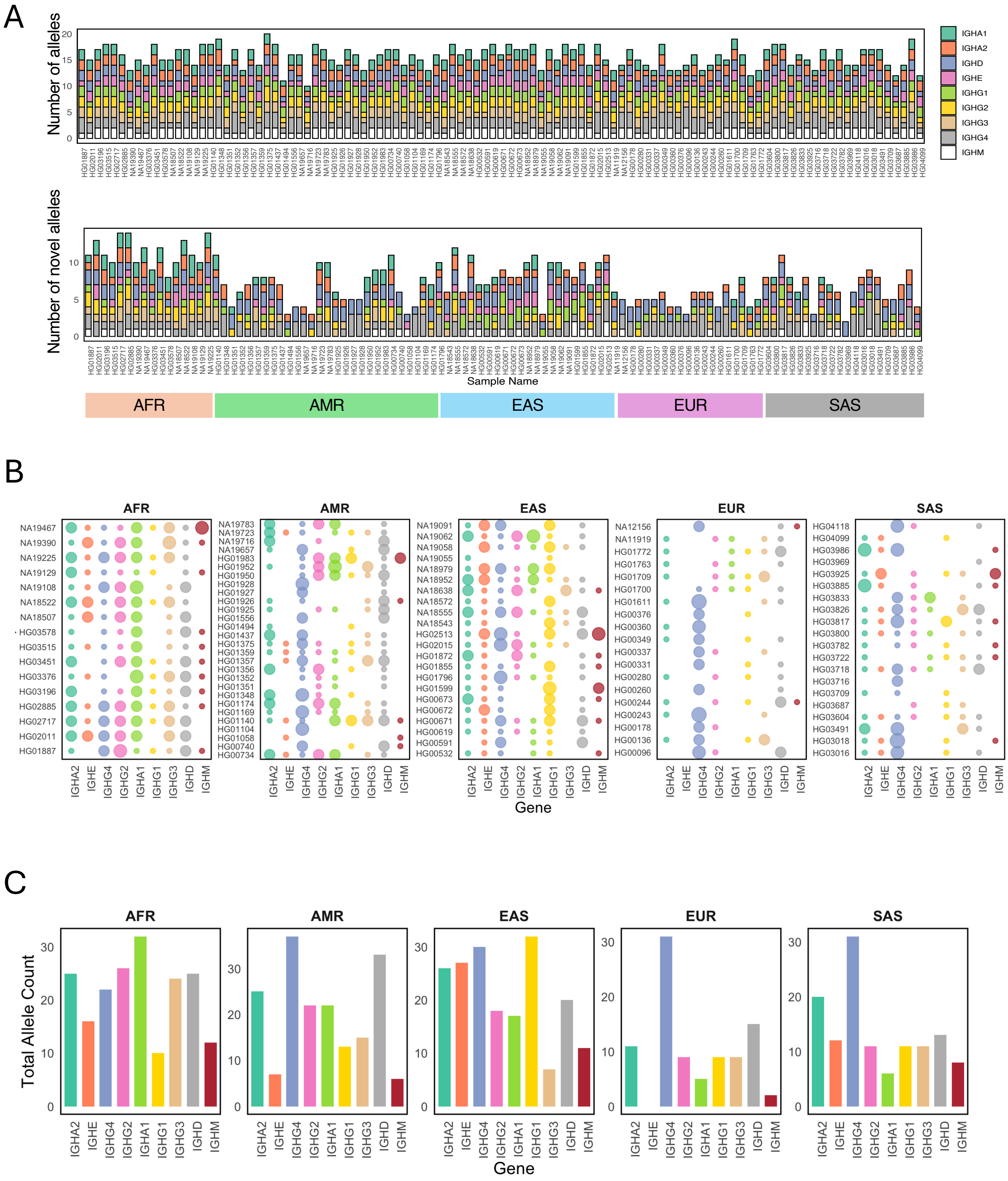


**Figure S10. Distribution of novel alleles over IGHC genes in individuals across diverse super-populations, related to Figure 5**

**(A-B)** Number of novel alleles counted across genes per individual from different super-population depicted by the stacked bar plot (A) and the bubble plot (B). **(C)** Number of novel alleles counted across genes grouped by the super population.

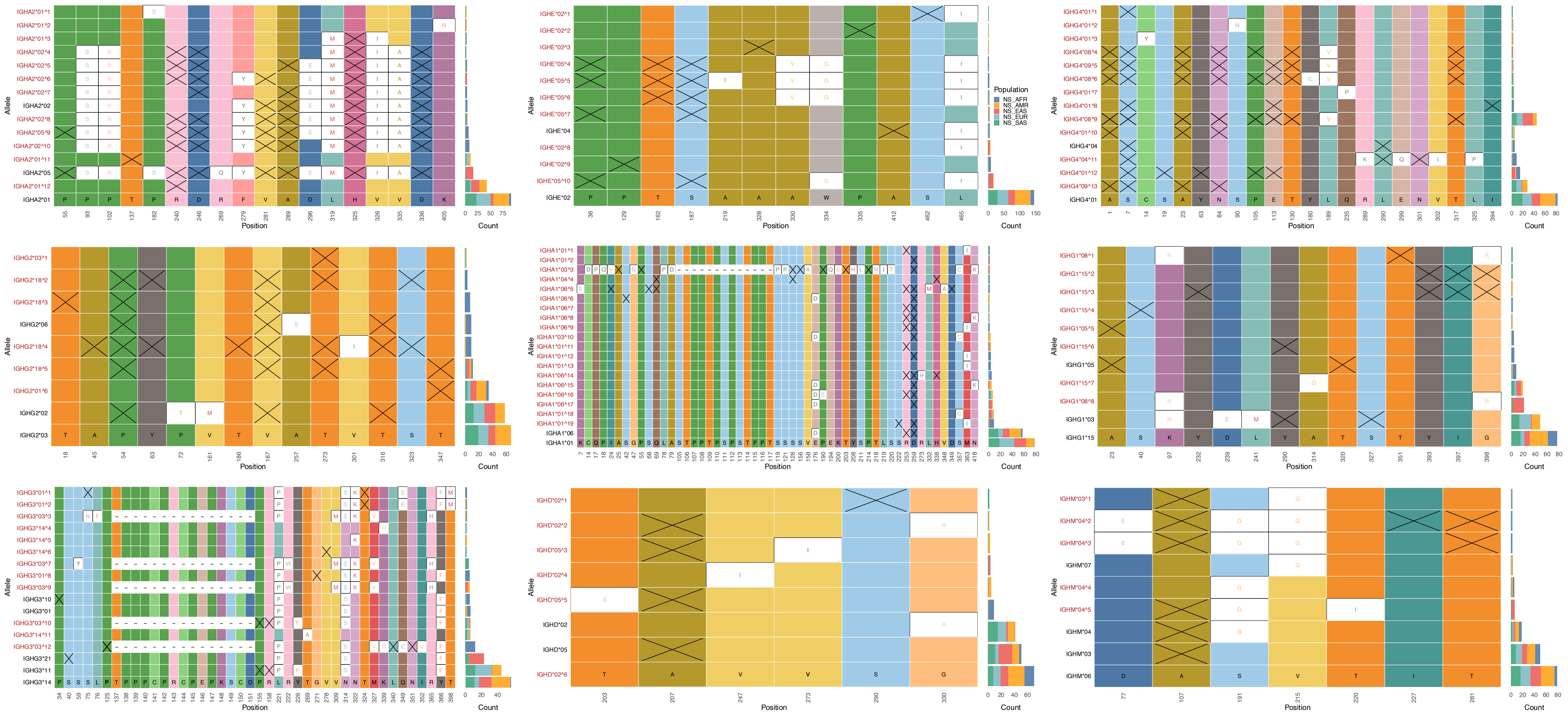


**Figure S11. SNV analysis reporting synonymous, non-synonymous changes compared to most frequent allelic variety as observed in all IGHC alleles (present in > 1 individual), related to Figure 5**

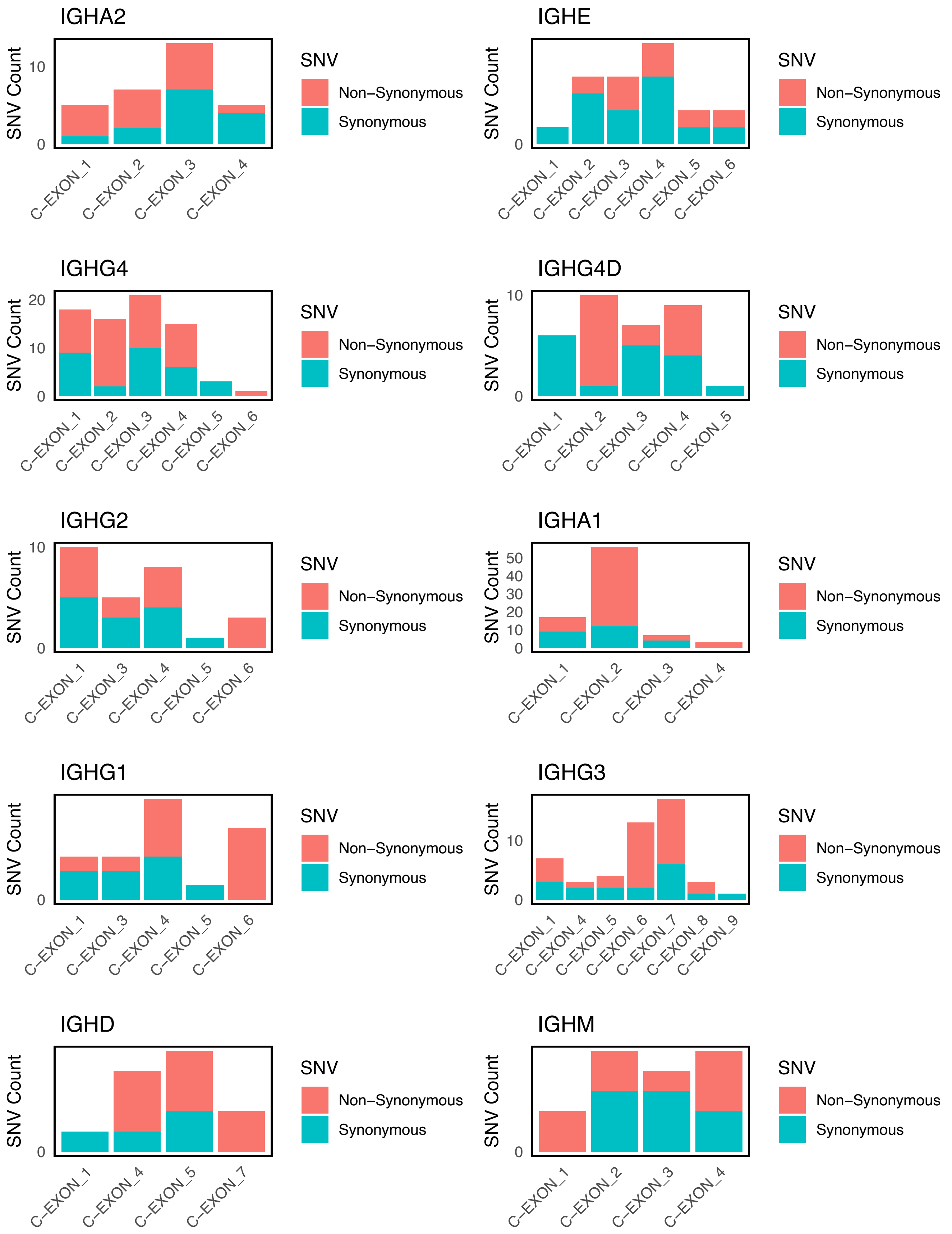


**Figure S12. Comparison of synonymous and non-synonymous changes in alleles compared to reference allele (based on ighc reference haplotype), related to Figure 4-5**

The IGHC alleles show a greater tendency for non-synonymous changes in the IGHA, IGHG, and IGHD isotypes compared to the reference haplotype, indicating higher genetic diversity in these genes. In contrast, IGHE alleles predominantly accumulate synonymous mutations, suggesting strong functional conservation. The IGHM genes exhibit a roughly equal distribution of synonymous and non-synonymous changes, pointing to a balanced evolutionary pattern

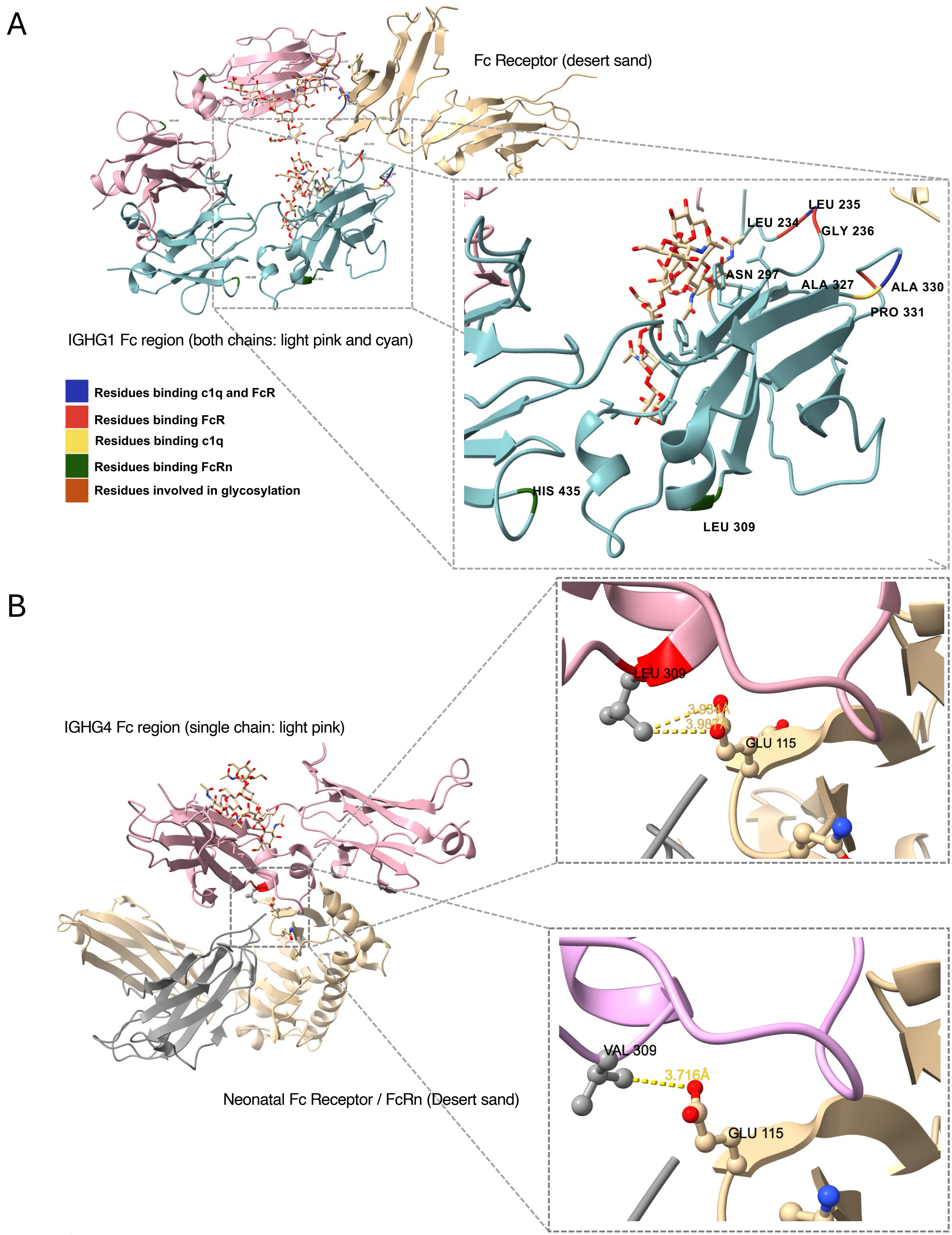


**Figure S13. Isotype specific lookup for non-synonymous mediated functional changes in IGHG alleles, related to Figure 4-5**

**(A)** Characterization of functionally important residues in typical IgG antibodies based on IGHG1 (pdb id. 1E4K shows Fc-FcR interaction **(B)** Isotype specific amino acids were preserved for the core functional residues, i.e., res. 234-L(IgG1,IgG3), V(IgG2), F(IgG4). Contradictorily, in IgG4 res. 309 harbored variation L309 like IgG1 and V309 like IgG2 possibly altering Fc-FcRn interactions.

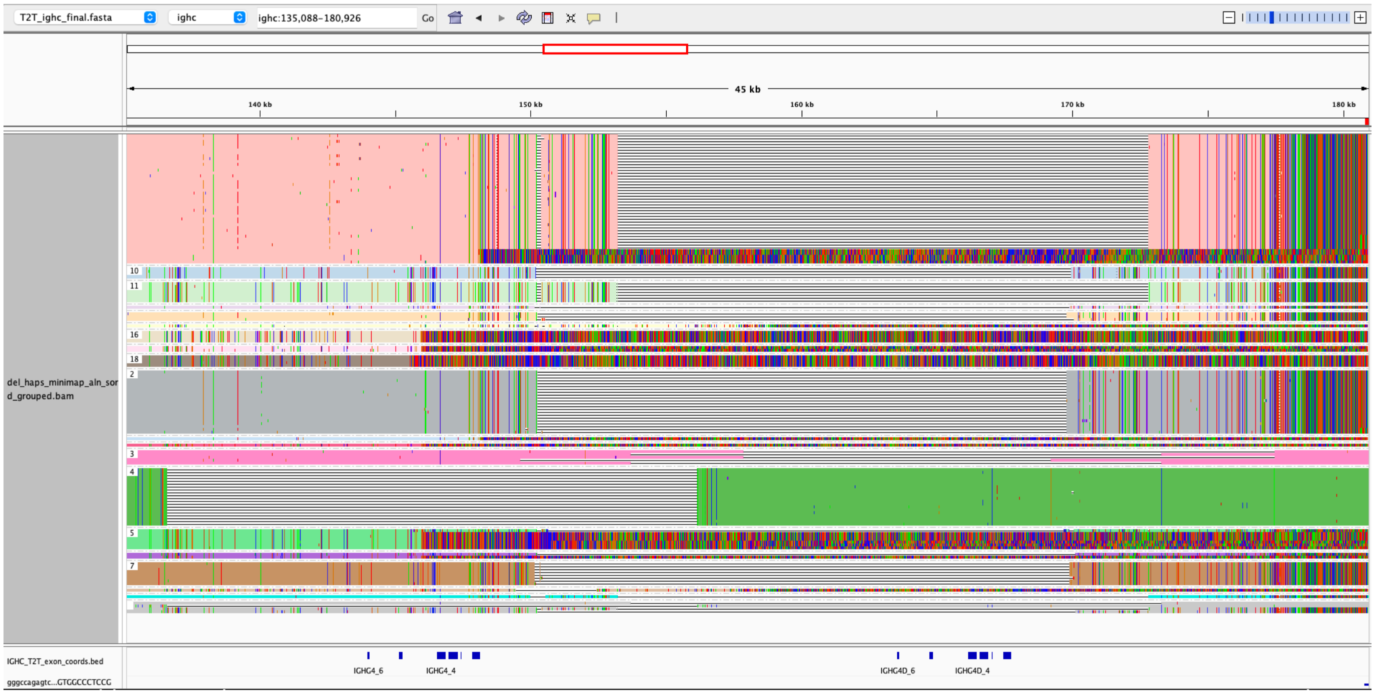


Different breakpoints observed in IGHG4 deletion haplotypes

Alternate genotyping approach for the complex SV

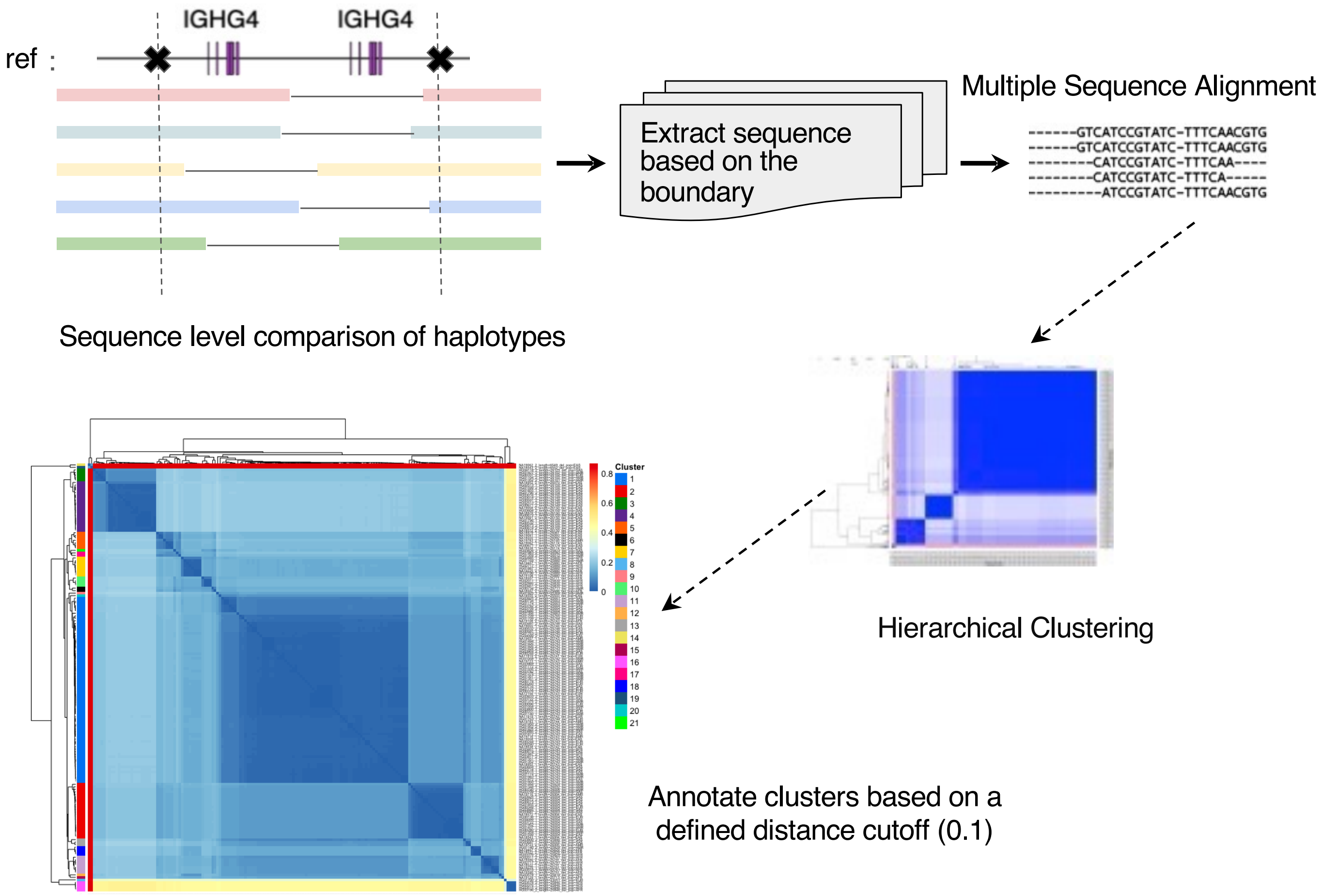


**Figure S14. Schematic representation of Genotyping approach for the complex IGHG4 deletion SVs, related to Figure 6 and STAR Methods**

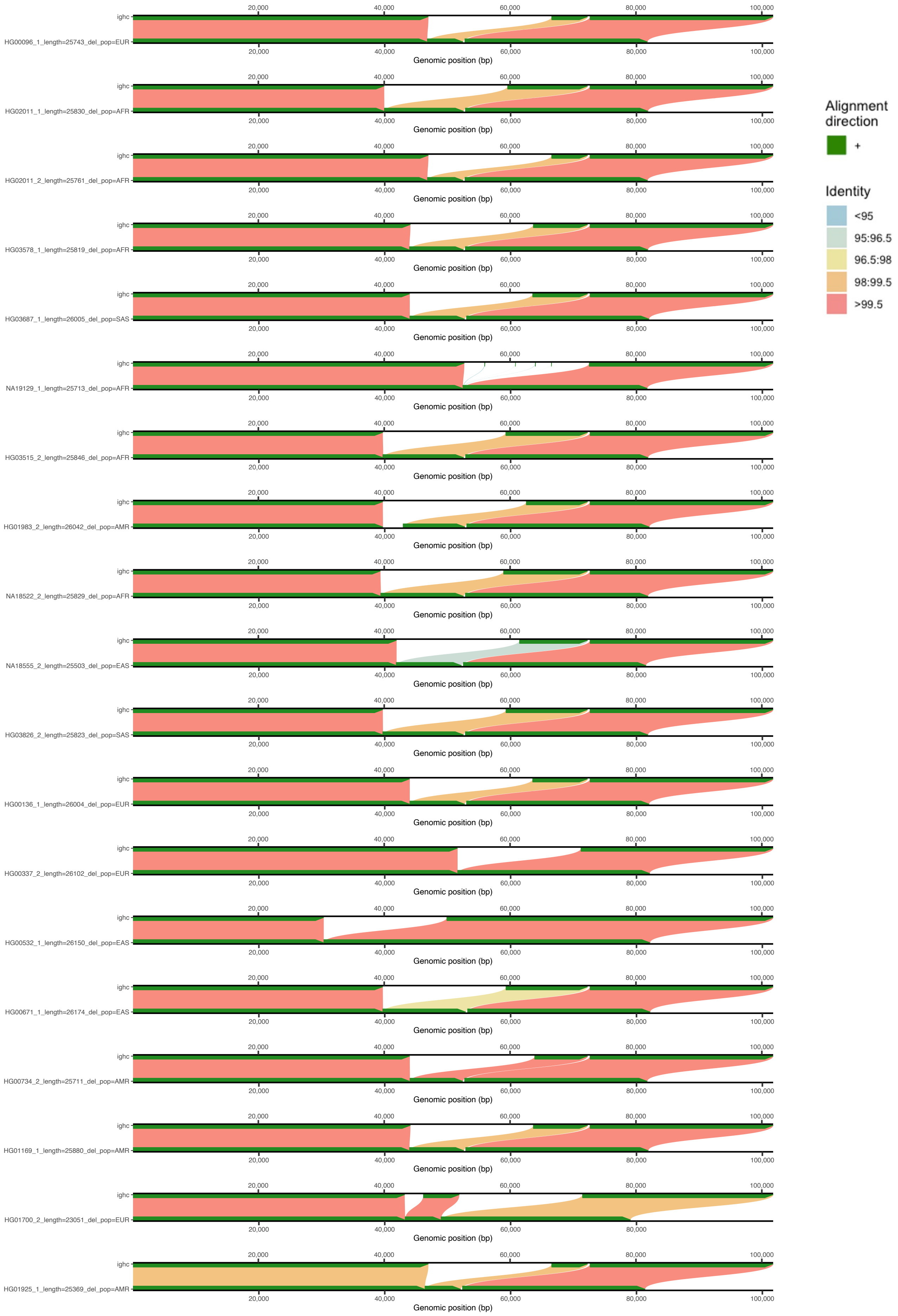


**Figure S14. (cont.) Schematic representation of Genotyping approach for the complex IGHG4 deletion SVs, related to Figure 6**

Here is a representation of the haplotype clusters obtained based on distance-based genotyping, showing how the deletion breakpoints compare with each other in terms of alignment over the specific IGHC region, using the CHM13v2.0 reference and analyzed with SVbyEye (Porubsky, David et al) [S1].

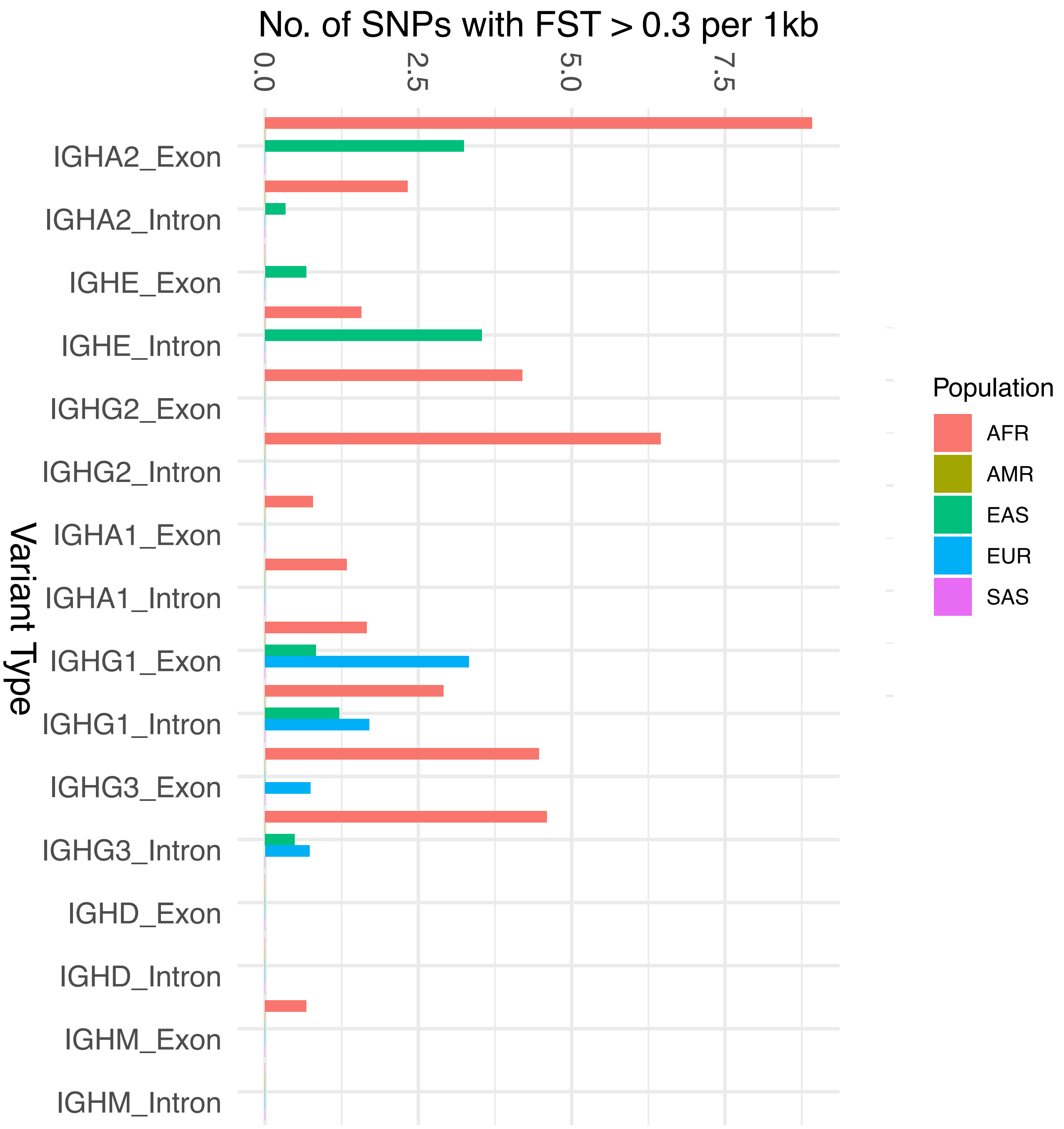


**Figure S15. Bar plot showing distribution of SNVs in the IGHC locus with high $F_{ST}$ values across different super-population, related to Figure 7**

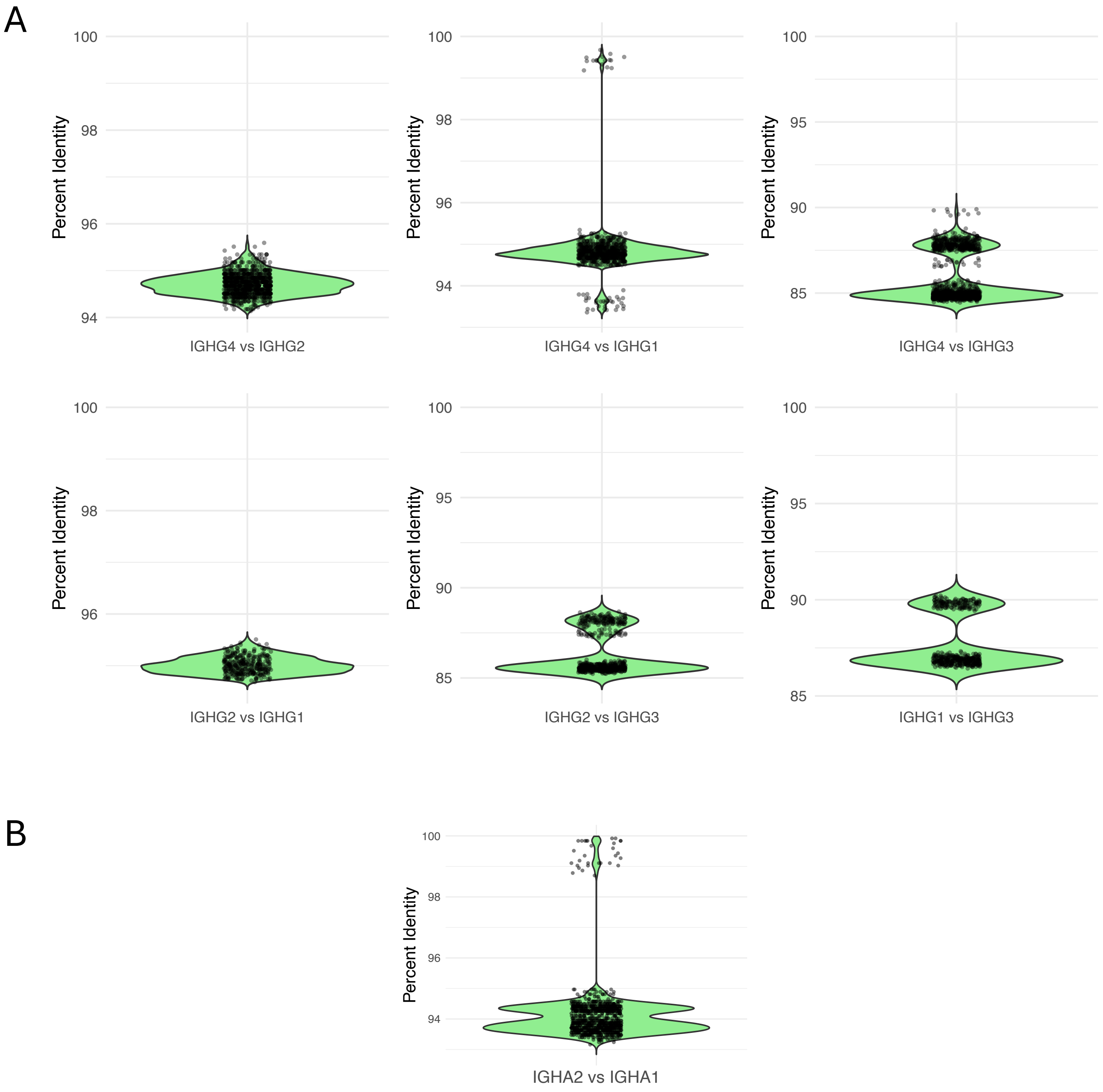


**Figure S16. Comparison of sequence identity among isotype defining paralogous genes: IGHG (G1, G2, G3, G4) and IGHA (A1, A2), related to STAR Methods**

**(A)** Violin plot illustrating the pairwise sequence identity among alleles of different IGHG genes. Among these, IGHG1, IGHG2, and IGHG4 alleles show the highest sequence similarity, indicating close evolutionary relationships. **(B)** Violin plot depicting pairwise sequence identity between all IGHA1 and IGHA2 alleles. The comparison reveals an average sequence identity of approximately 94%, suggesting strong conservation within the IGHA gene family.

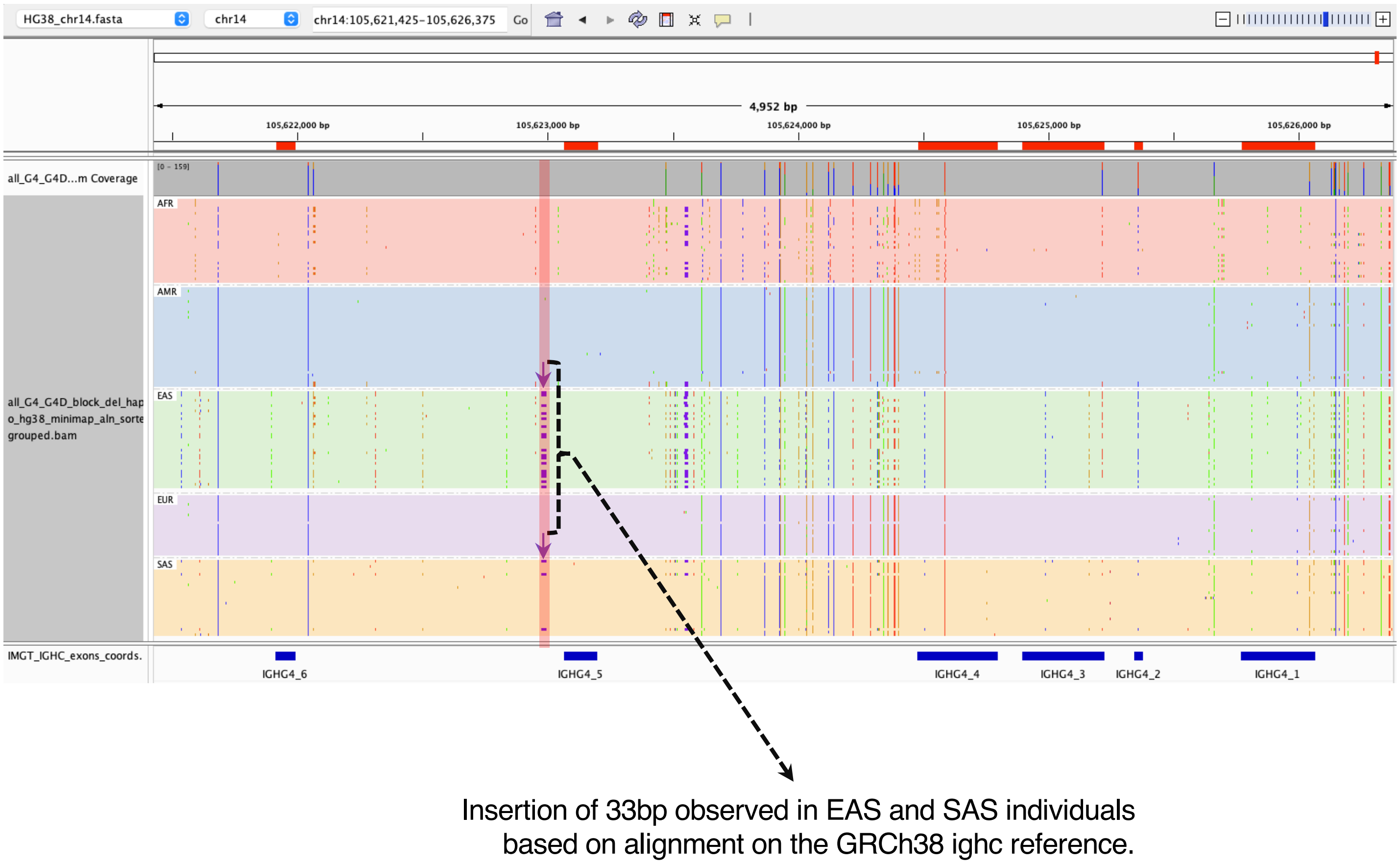


**Figure S17. Observation of ancestry defined evolutionary signature in the IGHC locus, related to Figure 7**
The ancestry-partitioned alignment of haplotypes over the IGHC locus in the GRCh38 reference suggests the presence of a 33bp nucleotide insertion signature in the intronic region of IGHG4, which was previously reported as evidence of Neanderthal introgression by Yan et al. (2021) [S2]. Our study aligns with their findings and further supports the higher ancestral bias for this insertion among haplotypes from the EAS and SAS super-populations.

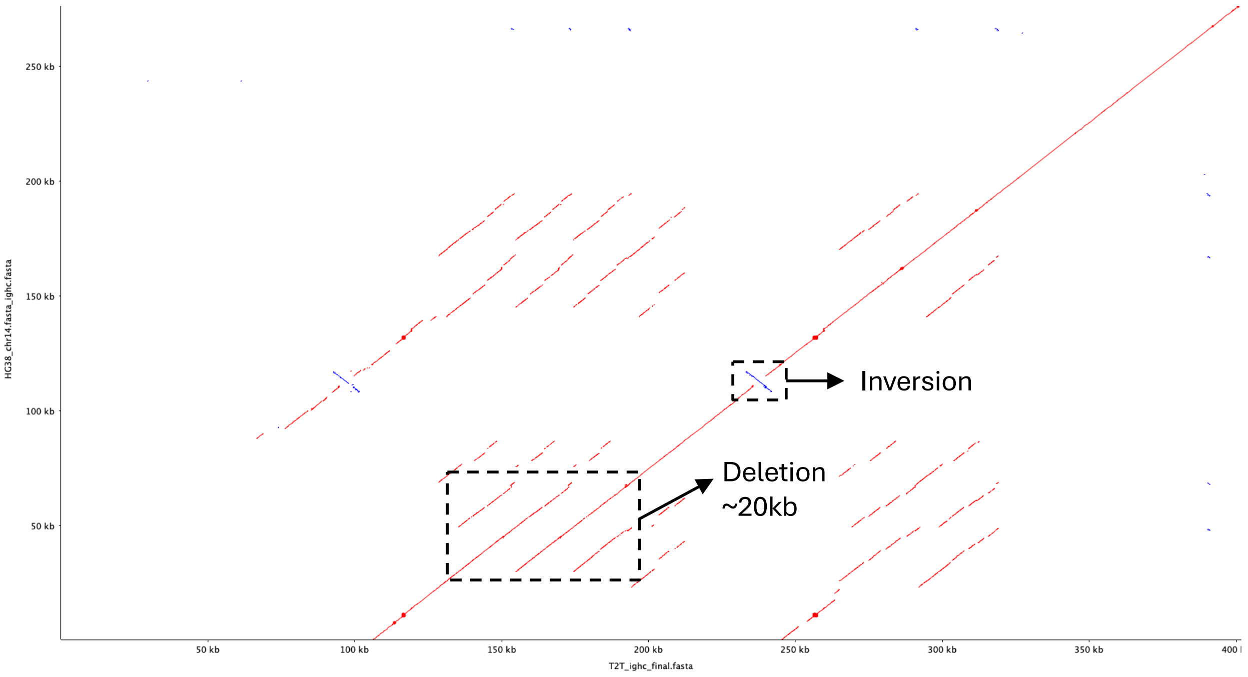


**Figure S18. A dot plot based comparison between the IGHC loci obtained from GRCh38 (X axis) and T2TCHM13v2.0 (Y axis) reference assembly, related to STAR Methods**
